# Deletion of major shell proteins of ethanolamine utilization microcompartment reduces intrinsic antibiotic resistance, biofilm, and intracellular survival of *Salmonella* Typhimurium

**DOI:** 10.1101/2025.07.17.665469

**Authors:** Minal R. Bhalerao, Aishwarya S. Davkhar, Ajit R. Sawant, Anindya S. Ghosh, Tiffany N. Harris, Thomas A. Bobik, Ankita Saha, Sachin B. Agawane, Chiranjit Chowdhury

## Abstract

With the high rise in *Salmonella* infection and emergence of antibiotic-resistant variants, developing a novel strategy to control the pathogen is imperative. Earlier studies revealed that *Salmonella* deploys ethanolamine (EA) metabolic machinery to disseminate in the intestine. *Salmonella* with a defect in EA metabolism manifests with lower intestinal colonization efficiency. Remarkably, the potential of EA metabolism as a therapeutic target is yet to explore. Our study revealed that supplementation of EA and vitamin B_12_ in both rich and minimal media enhanced biofilm formation, increased motility, and increased tolerance of *Salmonella* to some antibiotics. Conversely, mutants deficient in EA metabolic enzymes exhibited no physiological fitness. In *Salmonella*, EA metabolic enzymes are localized within a proteinaceous microcompartment (MCP) shell composed of thousands of copies of shell proteins encoded by five genes from the *eut* operon. Fascinatingly, bacterial cells with defective MCP shell due to mutation in the major shell proteins showed enhanced susceptibility towards a number of antibiotics in minimal media. The mutants were unable to form biofilm, produced lower curli expression and were defective in flagellar motility. Also, mutation in one of the major shell proteins reduced intramacrophagic viability of *Salmonella*. Notably, phenotypes were restored upon ectopic expression of corresponding genes. It was evident that mutation in the MCP shell proteins downregulated the expression of genes related to pathogenicity. Overall, this study sheds new light on understanding the relationship between EA metabolism and bacterial physiology that would pave the way for developing novel therapeutic interventions against *Salmonella*.

## 1. INTRODUCTION

Cases of gastroenteritis caused by non-typhoidal *Salmonella* (NTS) infection and subsequent deaths are on the rise (1, 2) due to the acquisition of multiple antibiotic resistance, which has made NTS refractory to commonly used antibiotics [1–9]. This opportunistic pathogen has an extraordinary capacity to establish colonization. It can form biofilms [10–17], which are advantageous for ensuring its survival in many ecological niches, rendering antibiotic treatment often unsuccessful. Hence, a new strategy of infection control is imperative.

Bacterial nutrient metabolisms have long been proposed as therapeutic targets [18, 19]. A recent study suggested that the metabolism selectively responsible for in vivo growth and dissemination of the pathogen can represent a new antimicrobial target [7, 18, 20]. It is indicated that EA metabolism bestows *Salmonella* with a selective proliferation advantage in aerobic and anaerobic environments [21–24] over natural gut bacteria that primarily do not host this function [25–28].

EA is a breakdown product of phosphatidylethanolamine, a phospholipid found in the intestinal epithelium [26]. Shedding of enterocytes from the intestinal epithelial cells and the host diets provides rich sources of ethanolamine in the gut environment [26]. It is evident that EA metabolism is essential for *Salmonella* fitness in the inflamed intestine, and a mutant defective in EA metabolism was documented with fewer CFU of the pathogen in the spleen, liver, and intestine of mice [22]. EA utilization also contributes to *Salmonella* disseminating from the intestines to the host tissue, driving further systemic infection [15, 29]. Hence, as several studies advocate, disruption of EA metabolism could offer a novel infection control measure to combat *Salmonella* infection without dismaying commensals [30–32]. Remarkably, the potential of EA metabolism for future therapeutic development is far less investigated.

Previously, a few gene expression studies with other bacteria claimed a link between EA metabolism and cellular motility and biofilm formation [14, 33–36]. It is proposed that biofilm formation reflects the potential adhering capability to intestinal surfaces [14, 37] and contributes significantly to pathogenicity and antibiotic resistance [12, 38]. EA metabolism may be *Salmonella*’s Achilles’ heel; however, direct evidence of EA metabolism on bacterial physiology remains elusive.

In *Salmonella*, the pathway of EA degradation begins with its conversion to ammonia and acetaldehyde by coenzyme-B_12_-dependent ethanolamine ammonia lyase (EAL) encoded by *eutB* and *C* genes [22, 25]. Ammonia is utilized as fixed nitrogen for purine and amino acid biosynthesis [39]; whereas, acetaldehyde undergoes further processing by acetaldehyde dehydrogenase (EutE) to convert it to acetyl-CoA [25](Fig. 1). Acetyl-CoA joins the central metabolism such as TCA cycle (energy generation), glyoxylate bypass (carbohydrate synthesis), and lipid biosynthesis. Alternatively, acetaldehyde may be converted to ethanol by alcohol dehydrogenase to maintain the cofactor pool up and running [25, 26] (Fig. 1).

**Figure 1.**
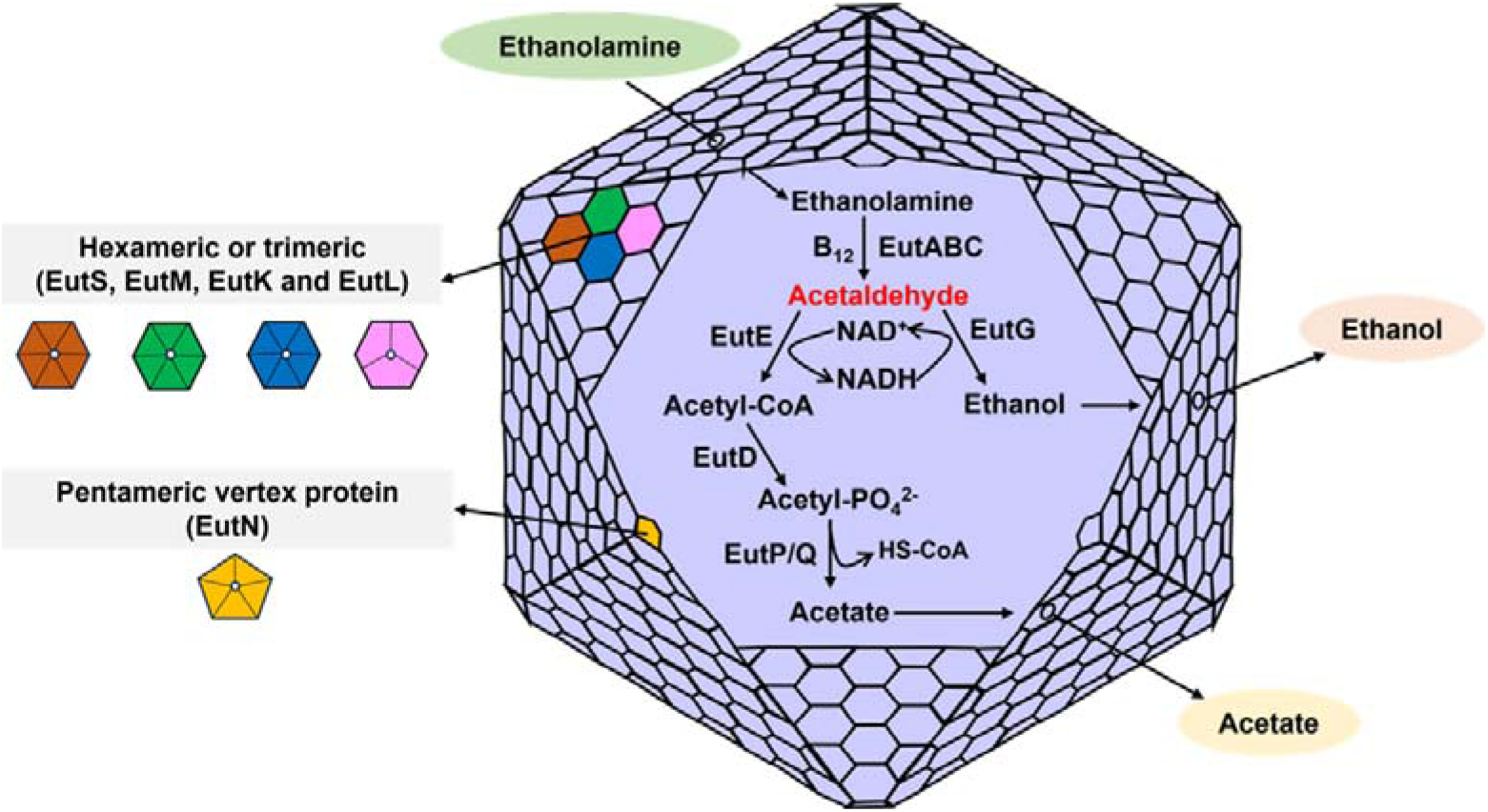
Functional model of Ethanolamine Utilization Microcompartment (Eut MCP).

It has been revealed that *Salmonella* encapsulates enzymatic pathways related to EA metabolism within a protein shell, known as the ethanolamine utilization microcompartment (Eut MCP) (Fig. 1) [23, 25, 26, 40–43]. Eut MCP creates an optimum environment for EA metabolism and provides the bacteria a unique advantage to survive in a hostile environment in the gut [44]. The shell of Eut MCP comprises five shell proteins, namely EutS, M, N, L, K [41, 45] that tessellate into facets, edges and vertices of the MCP shell [41, 46, 47] (Fig. 1). Genetic studies reveal the importance of shell proteins in proper MCP assembly [48–51], in retaining metabolic intermediate while selectively controlling the metabolic flux [52], and in the recruitment of enzyme cargo [43, 51, 53]. Earlier, deletion of the entire MCP shell has been demonstrated to abolish altogether EA metabolism in *Salmonella* (49), a phenotype similar to *eutBC* and *eutE* mutants that inactivate key metabolic enzymes (44, 51, 55), indicating a pivotal role of the MCP shell in EA metabolism. Recently, individual deletion mutants of *eutM*, *N*, *L*, or *eutK* shell protein of *Salmonella* were shown to impair MCP assembly, resulting in aberrant MCP morphology and bacterial growth defect [51]. It is suggested that the EA metabolic function was significantly affected in the shell protein mutants [51].

It has been proposed that disrupting EA metabolism by targeting microcompartment shell assembly could serve as a template for future therapeutic design [31]. Herein, with the help of molecular genetic techniques, we constructed individual shell protein mutants in *Salmonella enterica* Typhimurium LT2. The impact of mutations on bacterial physiology, intrinsic antibiotic resistance, and intramacrophagic survival are demonstrated.

## We envisaged that this study would help devise better therapeutic strategies to mitigate *Salmonella* infections in the future

## 2. MATERIALS AND METHODS

### 2.1 Chemicals and reagents

Ethanolamine hydrochloride, Cyanocobalamin (Vit. B_12_), glycerol, 3-Methyl-2-BenzoThiazolinone Hydrazone hydrochloride (MBTH), and Congo red were from Sigma-Aldrich (St. Louis, MO). Antibiotics, crystal violet, Isopropyl-β-D-1-thiogalactopyranoside (IPTG), and arabinose were from Himedia (Mumbai, India). LIVE/DEAD BacLight Bacterial Viability Kit was from Invitrogen (Waltham, MA). Q5 DNA polymerase, restriction enzymes, and T4 ligase were from New England Biolabs (Beverly, MA). NovaTaq PCR Master Mix was from Merck (Darmstadt, Germany). Other chemicals were obtained from Qualigens Thermofisher Scientific (Powai, India).

### 2.2 Bacterial strains, media, and growth conditions

The bacterial strains used are listed in Table S1. All strains are derived from *Salmonella enterica* serovar Typhimurium strain LT2. The rich medium used was Luria-Bertani/Lennox (lysogeny broth) (LB) medium (Himedia, Mumbai, India). For selecting transformants TYE media was used, where bacto-tryptone, yeast extract and agar were purchased from Himedia Laboratories (Himedia, Mumbai, India). For MIC experiments, overnight cultures were inoculated in cation-adjusted Mueller–Hinton broth (MHB) and subcultured into MHB or the indicated minimal media. For biofilm experiments the rich media used was Tryptic Soy Broth (TSB, Himedia, Mumbai, India) [54]. The minimal media used were either no carbon-E (NCE) or no carbon-no nitrogen (NCN) medium [55]. Ethanolamine (EA) (30 mM) was used as a carbon source in NCE medium, as a nitrogen source in NCN medium with glycerol (20 mM), or as a carbon and nitrogen source in NCN medium [55]. Minimal media were supplemented with 1 mM MgSO4, 0.3% each of valine, isoleucine, leucine and threonine and 50 µM ferric citrate [49, 56]. CN-B_12_ or Vit. B_12_ (150 nM) was served as exogenous B_12_ source. Antibiotics were used at the following concentrations: ampicillin, 100 μg/mL and chloramphenicol, 20 μg/mL. Solid media were prepared by adding agar (1.5%, Himedia) to LB, TYE, NCE, or NCN medium.

### 2.3 Construction of deletion mutants and complemented Strains

All primers and plasmids used in this study are listed in Table S2. Chromosomal deletion of individual Eut MCP shell proteins was constructed by recombineering as follows [49, 57, 58]. In brief, the chloramphenicol cassette flanking the FRT sequence was amplified from the pKD3 plasmid so that the amplicon contains portions of sequences adjacent to the gene to be deleted. The amplicon was electroporated into the cell expressing λ-Red recombinase from plasmid pKD46. Transformants were screened against chloramphenicol (20 µg/mL) on TYE-Cam agar plates. The antibiotic cassette was crossed off by expressing flippase from the plasmid pCP20. The chloramphenicol cassette at respective gene loci was moved to another mutant by transduction (74, 75), using phage P22 HT105-int to make double mutants. For complementation studies, the genes encoding Eut shell proteins were cloned into the pLac22 plasmid within the *Bgl*II and *Hin*dIII sites as stated earlier [49]. All the clones and mutants were confirmed by PCR followed by sequencing.

### 2.4 Antibiotic susceptibility assays of various classes of antibiotics

Initially, the antibiotic sensitivity experiment was carried out in the agar diffusion assay (both MHB and minimal media supplemented with 30 mM EA and 150 nM B_12_) using EZY-MIC strips (Himedia) of structurally unrelated classes of antibiotics, such as ampicillin, cephalothin, chloramphenicol, ciprofloxacin, cefotaxime, kanamycin and piperacillin. The minimum inhibitory concentration (MIC) based on the Epsilometer method (E-test) was used to determine the antimicrobial sensitivity of the WT *Salmonella* and individual shell protein mutants. Later, the broth dilution method was adopted to enhance the test’s sensitivity. The concentration range of antibiotics for the broth dilution method was obtained from the agar diffusion assay. The micro-broth dilution assay was performed in 96-well microtiter plates according to the Clinical and Laboratory Standards Institute (CLSI) guidelines [59]. Overnight cultures were inoculated from a single colony into LB and subcultured 1:100 into the cation-adjusted MHB supplemented with 30 mM EA and 150 nM B_12_ and grown until OD_600_ reached to ∼1.0. Cultures were diluted into either cation-adjusted MHB or minimal media for a starting inoculum of ∼1 × 10^5^ CFU/mL. In order to prime the initial cell growth, the minimal media with EA and B_12_ was supplemented with 0.5 mM glycerol, as described elsewhere [48]. The antibiotics were twofold serially diluted, and the total volume was maintained to 200 μL. Assay plates were incubated for 24 h at 37 °C with double orbital shaking. The growth was monitored with a BioTek Epoch 2 Microplate Spectrophotometer (Agilent-BioTek, USA). The assay was replicated four times, and at least three biological replicates were used. The MIC was calculated based on the concentration that inhibited bacterial growth by >90% compared to the no antibiotic control. To determine intrinsic growth differences between WT and individual shell protein mutants, a relative antimicrobial susceptibility index (ASI) was calculated, which can be calculated as the ratio of (mutant OD_600_/wild-type OD_600_) in the presence of antibiotics/ (mutant OD_600_/wild-type OD_600_) without antibiotics. ASI values less than 1 indicate an increased antimicrobial susceptibility.

### 2.5 Determination of Biofilm formation by crystal violet assay

Biofilm formation assay was performed as described earlier with some modifications [60]. Overnight cultures of wild-type *Salmonella*, its mutants, and the cells harboring their respective plasmid clones were diluted 1:100 in TSB and were allowed to grow until OD600 reached 0.5. The cells were preinduced for microcompartment formation with ethanolamine (30 mM) and B_12_ (150 nM). The culture was further diluted into biofilm growth media (either TSB or minimal media supplemented with ethanolamine and B_12_) into 96-well polystyrene plates to an OD_600_ of 0.1, and incubated at 37 °C for 72 h under static conditions. The genes from the respective plasmid clones were expressed by inducing with 20 μM IPTG.

After incubating the plates for 72 h, the wells were washed twice with phosphate-buffered saline (PBS), and the biofilms were fixed at 80 °C for 15 min, followed by staining with 0.5 % crystal violet (CV) for 15 min. The excess stain was rinsed with distilled water followed by air-dried. The bound CV on the biofilms was solubilized by 33% acetic acid (v/v) and measured at OD_570_ using an Epoch microplate spectrophotometer (Agilent-BioTek, USA). The biofilm formation index was calculated as BFI = (AB-CW)/(GB-GW), where AB is OD_570_ of the stained test well; CW is OD_570_ of the stained control well; GB is OD_600_ of the planktonic cells; and GW is OD_600_ of the control well [61].

### 2.6 Microscopic analysis of biofilm structures

The CV-stained biofilm in the microplate was observed under a 40× objective of the Labomed TCM400 microscope. Bacterial viability within the biofilm structures was determined by staining the biofilm samples on a coverslip with LIVE/DEAD BacLight Bacterial Viability Kit (Invitrogen, MA) per the manufacturer’s instructions. After washing the biofilm with PBS, the stained samples were viewed under a confocal laser scanning microscope (Leica Stellaris, Germany) at 20× magnification. The images were captured and processed with the help of LASx software to determine the thickness of the biofilm matrix and the bacterial viability within its structure.

The biofilm architecture was also determined with the help of field-emission scanning electron microscopy (FE-SEM). Samples were prepared by washing the air-dried biofilm containing cover slips with PBS, followed by fixation with 2.5 % glutaraldehyde. Samples were gold-coated and finally subjected to visualization under a FEI Nova Nano 450 field emission scanning electron microscope and the images were captured at 5000× magnification.

### 2.7 Assessment of *Salmonella* morphotypes by Congo red agar plate assay

The curli morphotypes of wild-type *Salmonella* and its mutants were screened in Congo red agar (CRA) plate as stated earlier [62] with some modifications. Overnight bacterial cultures in LB were diluted to secondary culture media (LB supplemented with 30 mM ethanolamine and 150 nM cyanocobalamin) at a ratio of 1:100. Next, optical density (OD) of the cultures were adjusted to an OD_600_ ∼ 0.2 with phosphate buffered saline. The diluted culture was next spot inoculated (5 μL) onto either Tryptone-Yeast extract (No NaCl) agar plate or in the minimal media agar plate supplemented with EA and B_12_, as stated before. Congo red (40 mg/L) (Sigma-Aldrich, St. Louis, MO, USA) was added to the media as mentioned earlier [62]. The colony morphology and the color were monitored after 96 h of incubation at 37 °C.

### 2.8 Calcofluor assay for EPS production

Bacterial EPS production was assessed in a calcofluor agar plate, as discussed earlier with some modifications [62]. Overnight cultures of WT *Salmonella* and respective mutants were diluted to 1:100 in LB supplemented with 30 mM ethanolamine and 150 nM Vit B_12_ and were grown for six hours at 37 °C. The optical density of all the cultures was adjusted to ∼ 0.1 before streaking onto the calcofluor-agar plates supplemented with 1 mg/mL calcofluor white (HiMedia). Plates were incubated at 37 °C for 72 h, and the fluorescence intensity was captured under a UV light source. EPS production was also quantified by measuring the calcofluor bound cells as described previously [62]. Fluorescence measurements were performed in a Fluorescence Spectrophotometer (Promega) with excitation at 360 nm and emission at 460 nm. The fluorescent intensity in arbitrary units was normalized with the OD_600_ of the bacterial suspension and represented relative to WT. Data are mean values of triplicate determinations plotted with the standard error of the mean.

### 2.9 Motility Assay

For swarming motility behavior studies, the concentration of agar used in the media was 0.6% (w/v). Different combinations of minimal medium were used, where ethanolamine served as a carbon or nitrogen source or both. Overnight cultures of WT *Salmonella* and respective mutants were diluted to 1:100 in LB supplemented with 30 mM ethanolamine and 150 nM Vit. B_12_ and were grown for six hours at 37 °C. One microliter of the culture with OD_600_ ∼ 0.1 was spotted onto the minimal medium agar plates. The swarming pattern was observed after a 72 h incubation period and the diameter was measured.

### 2.10 RNA extraction and qPCR analysis

Total RNA was extracted from biofilm-dwelling bacterial cells and the cells from the swarming agar plates. Both planktonic and sessile cells from biofilm plates and cells from swarming plates were resuspended in 500 µL PBS. Double volume of RNAprotect bacterial reagent (Qiagen, Hilden, Germany) was added to stabilize RNA before isolation. RNA was extracted using the RNeasy Mini Kit (Qiagen, Hilden, Germany) according to the manufacturer’s instructions. RNA purity was determined by measuring the A_260_/A_280_ absorbance ratio, and the concentration was determined with a SimpliNano Spectrophotometer (GE Health Care, UK). RNA was converted to cDNA using Ecodry Premix (Random Hexamers, Takara, Japan). The cDNA was amplified using the Applied Biosystems 2x SybrGreen Master Mix as per manufacturer’s protocol by consensus primers detecting the expression genes related to biofilm, motility and cell invasion, as stated in Table S2. Primers were designed using Primer 3, GeneRunner, and Multiple Primer Analyzer (Thermo Fisher Scientific) to ensure no cross-reactivity to other genes in the *Salmonella* LT2 chromosome. Amplicon length was approximately 150-200 bp. Reactions were run using the StepOnePlus RT-qPCR machine (Applied Biosystems, USA). Three technical replicates were averaged for analysis by the relative quantification method in which C_T_ values were normalized to the reference genes *rpoA* and *gyrA.* The expression level of the target genes was compared using the relative quantification method. Data were presented as the change (n-fold) in expression levels of the target genes in mutants as compared to that in WT. Error bars represent the standard errors with mean of the 2^-ΔΔCT^ values.

### 2.11 Estimation of acetaldehyde

The wild type and Eut shell protein mutants were grown in 1 mL of minimal medium containing 30 mM ethanolamine and 150 nM of Vit. B_12_ for acetaldehyde measurement. The culture was incubated in 5 mL sterile glass vials containing rubber caps that were crimp sealed with aluminum caps with the help of a crimper tool. The glass vials were incubated at 37 °C for 12 h. A 48-well microplate (VWR) was prefilled with 286 µL of 100 mM Potassium citrate buffer (pH 3.6), 143 µL of 0.1% MBTH and 246 µL of double deionized water. Culture sample of 40 µL was collected from each vial using a sterile syringe without opening the cap and was subjected to add the respective well. The plate was incubated at 37 °C for 15 minutes under static conditions. After incubation, the samples in the wells were diluted by adding 285 µL of double-deionized water. The absorbance of the samples was measured at 305 nm in the microplate reader as described previously [63].

### 2.12 Bacterial infection in macrophages

The RAW264.7 murine macrophage cell lines were grown and maintained in Dulbecco’s Modified Eagle Medium (DMEM) with 10 % Fetal Bovine Serum and penicillin-streptomycin solution at 37 °C in 5 % carbon dioxide. The macrophage cells were grown in modified DMEM media supplemented with FBS (10%), ethanolamine (30 mM), Vit. B_12_ (150 nm), and Sodium pyruvate (1 mM) for the infection experiment. Overnight cultures of both wild-type and mutants of *Salmonella* in LB broth were diluted 1:100 in fresh LB media supplemented with ethanolamine and vitamin B_12_ and were allowed to grow until the mid-log phase. The bacterial cells were harvested and diluted to an OD_600_ of 0.1. RAW264.7 macrophages were infected with these bacteria at a multiplicity of infection (MOI) of 50:1 and centrifuged at 1000 rpm for 5 min to aid the adhesion of bacterial cells to the macrophage. The plates were kept undisturbed for 60 min at 37 °C in 5 % carbon dioxide.

### 2.13 Microscopic analysis of bacterial viability within infected macrophages

To visualize bacterial cells inside the macrophage RAW264.7, the wild-type and mutants were transformed with a plasmid (pLac22-mCherry) expressing mCherry (Red fluorescence) after the addition of IPTG (1 mM). The infection of the macrophage was done as explained in the above section. One hour after infection of the RAW264.7 cells with *Salmonella* strains, the media were replaced with gentamycin (100 µg/mL) containing DMEM media to kill the non-internalized bacterial cells and incubated for one hour. Further, the gentamicin media were replaced with DMEM media without antibiotics and incubated for two hours. The macrophage cells were washed gently with PBS to remove any non-internalized bacteria before being subjected to imaging. The fluorescence microscopic images of the viable bacteria within macrophages were captured using a fluorescence microscope with the Excitation/Emission for mCherry: 587/610 nm.

## 3. RESULTS

### 3.1 Deletion of shell proteins makes *Salmonella* susceptible to structurally unrelated antibiotics

Although minimum inhibitory concentrations (MICs) of some antibiotics had been lower in some shell protein mutants (Table S3), changes in the susceptibilities to various antibiotics attributed to shell protein deletion mutation could not be explained by changes in the MICs. Earlier, it has been suggested that the effect of mutation on bacterial growth in the presence of an antibiotic as compared to the wild-type could be better analyzed by determining a relative antibiotic susceptibility index (ASI) [64, 65]. To assess intrinsic growth differences between wild-type (WT) *Salmonella enterica* serovar Typhimurium LT2 and its shell protein mutants, a comparative ASI was calculated as the ratio of (mutant OD_600_/WT OD_600_) with antibiotics/(mutant OD_600_/WT OD_600_) without antibiotics, as suggested previously [64, 65]. Here, WT *Salmonella* and its shell protein mutants were tested with a variety of antibiotics, viz, ampicillin, cephalothin, ciprofloxacin, kanamycin, chloramphenicol, cefotaxime, and piperacillin. The susceptibility index less than 1 indicates an increased susceptibility against a particular antibiotic (Fig. 2). Notably, the effect of antibiotics on the growth of shell protein mutants was evident in the No-carbon E minimal media supplemented with Vit. B_12_ and EA (NE media), not in the rich media like Mueller Hinton (MH) media. It has been observed that loss of shell proteins significantly increased the susceptibility of individual mutants to some of the antibiotics (Fig. 2, Fig. S1, S2). Out of all individual shell protein mutants, an *eutL* mutant became sensitive to the maximum number of antibiotics tested (Fig. 2, Fig. S1). In contrast, an *eutS* mutant behaved similarly to WT. Again among all the antibiotics tested, only ciprofloxacin and kanamycin showed activity to almost all the shell protein mutants (Fig. 2). Next, to substantiate the relationship between EA metabolism and intrinsic antibiotic resistance, the same test was conducted with *eutB* and *eutKM* mutants against ciprofloxacin. WT *Salmonella* presented more tolerance to ciprofloxacin in the presence of media containing EA and Vit B_12_ than in media containing glycerol alone (Fig. S3). Conversely, an *eutB* mutant with loss of EA metabolism failed to show any fitness even in the presence of EA and Vit. B_12_ (Fig. S3), confirming the role of EA metabolism in conferring intrinsic antibiotic resistance. Meanwhile, an *eutKM* mutant, where both EutK and EutM shell proteins were deleted, also behaved similarly to the *eutB* mutant (Fig. S3). The *eutKM* mutant did not grow in NE media (Fig. S4), presumably due to the complete disintegration of the Eut MCP assembly, which inhibited the bacteria from utilizing ethanolamine. These studies indicated that disruption of the Eut MCP shell reduced the intrinsic antibiotic resistance of *Salmonella* Typhimurium.

**Figure 2.**
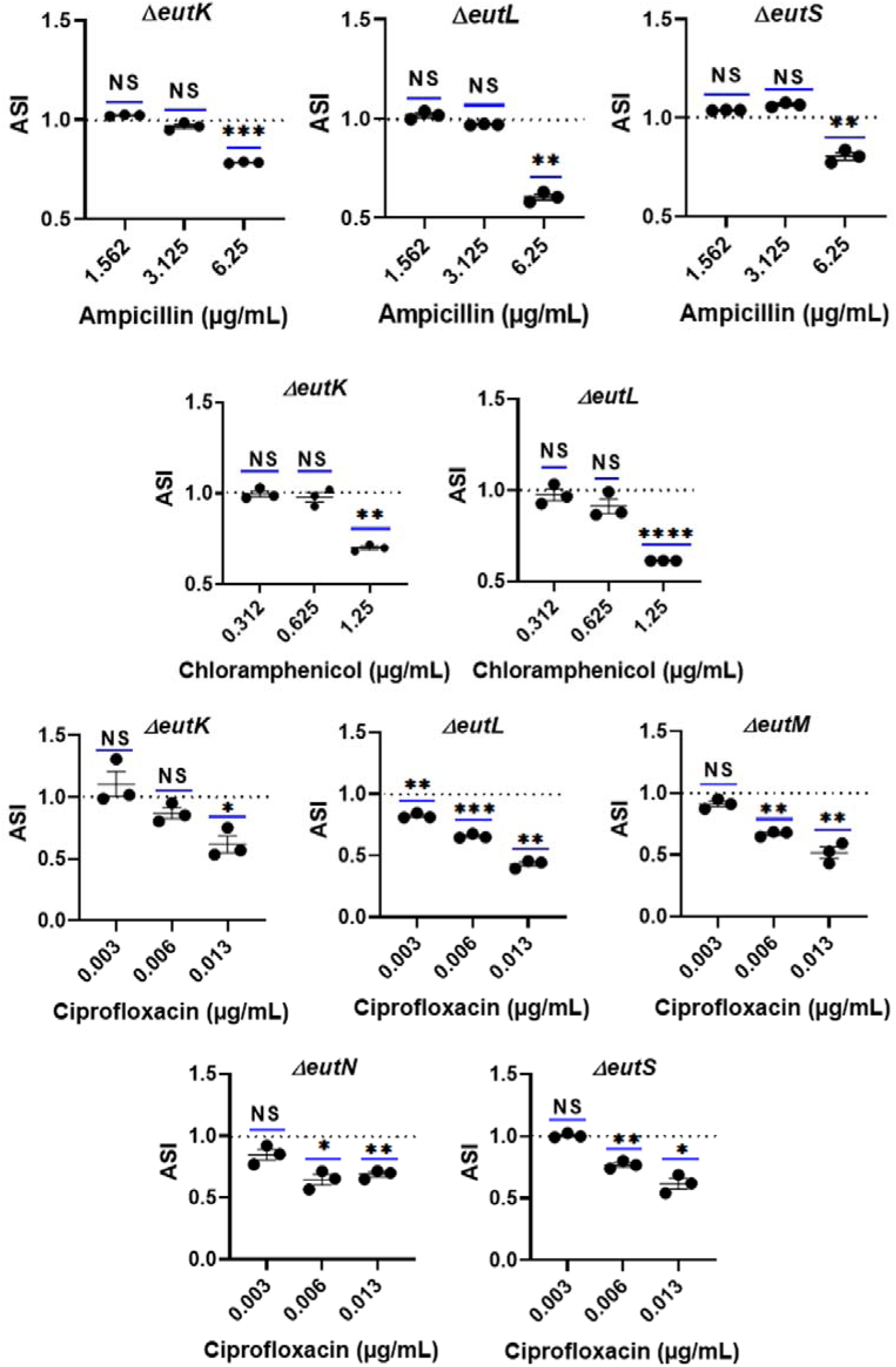
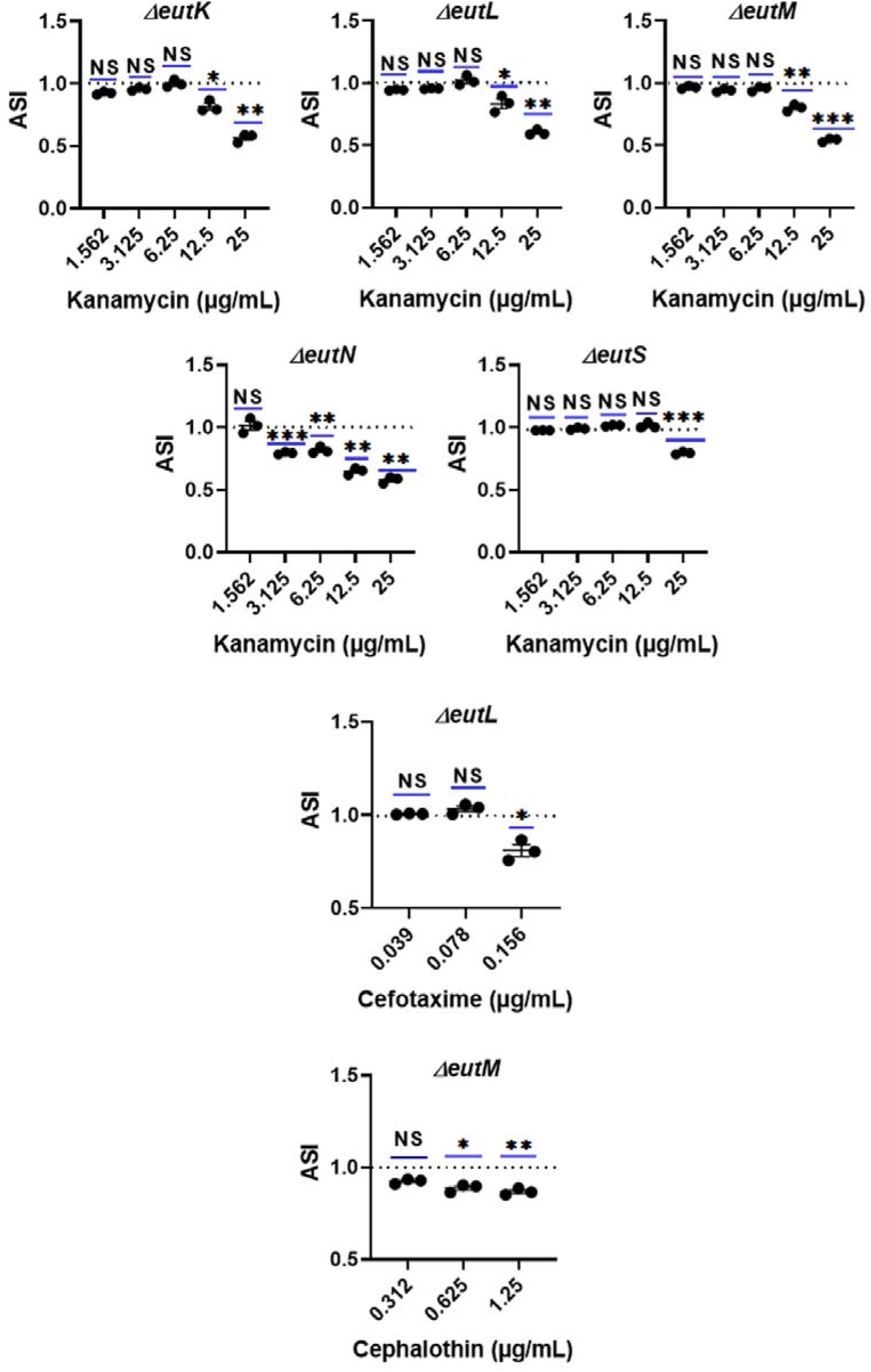
Antibiotic Susceptibility of *S*. Typhimurium (WT) and Eut shell protein mutants in the presence of different class of antibiotics. Growth of WT and mutant strains in minimal medium supplemented with 30 mM ethanolamine, 150 nM Vit. B_12_, and 0.5 mM glycerol was monitored at 37°C for 24 h by measuring optical density at 600 nm to evaluate the activity of the selected antibiotics. Three biological replicates and four technical replicates were performed for each antibiotic. The antibiotic susceptibility index (ASI) is the ratio of (mutant OD_600_ /wild-type OD_600_) with antibiotic/ (mutant OD_600_ /wild-type OD_600_) without antibiotic. ASI values less than 1 indicate an increased antibiotic susceptibility for mutants. NS p > 0.05, *p ≤ 0.05, **p ≤ 0.01, ***p ≤0.001, ****p ≤ 0.0001 (one-sample t test). The mutants where ASI values are similar to 1 have been shown in the supplemental file.

### 3.2 EA metabolism is impaired in the shell protein mutants

The microcompartment shell is known to retain the metabolic intermediates and channelize them to the downstream enzymes to keep the metabolic process up and running. Any perturbation in the retention of intermediate results in defects in metabolism and subsequently affects onto the growth. Earlier deletion of individual shell protein breaks Eut MCP assembly and impairs growth of *Salmonella* in the minimal media supplemented with EA and Vit. B_12_ [51]. In our study, we observed that the most shell protein mutants had poor retention of metabolic intermediate within Eut MCP, which gave rise to releasing more aldehyde in the media, except *eutK* and *eutS* mutants (Fig. S5). An *eutL* mutant released the highest acetaldehyde among all the shell protein mutants (Fig. S5); which indicated that the EA metabolism was greatly affected upon loss of EutL. It also suggested the major contribution of EutL shell protein in MCP assembly and function (Fig. S5).

### 3.3 Shell protein mutants have reduced the biofilm-forming ability of *Salmonella*

The biofilm-forming ability was assessed to determine whether the reduction of intrinsic antibiotic resistance to various antibiotics was likely to be a function of the impact of individual shell protein mutations on biofilm formation.

The biofilm was assessed quantitatively by the crystal violet assay and live/dead staining, followed by its visualization under confocal laser scanning microscope. Supplementation of EA and Vit. B_12_ in both Tryptic Soy Broth (TSB) and No-carbon E-glycerol (NG) media enhanced biofilm of wild type *Salmonella* as observed by crystal violet staining assay (Fig. S6). Notwithstanding the visual evidence, the phenotype was subtle in rich media and was quantitatively indistinguishable (Fig. S6a). Conversely, the phenotype was apparent when minimal media was used. The biofilm of wild-type *Salmonella* was observed to be enhanced by ∼20% when EA and Vit. B_12_ was supplemented in NG (NGE media) (Fig. S6b, c). Conversely, adding the same did not impart much effect in the enzyme mutant where EA metabolism was abolished entirely (Fig. S6b, c), indicating a relationship between EA utilization and biofilm formation. Interestingly, a *eutKM* mutant did not show any impact, even when EA and Vit. B_12_ was added (Fig. S6b, c).

Next, we set out to investigate the effect of an individual shell protein mutant on biofilm formation. It was observed that the phenotype was more discernible in the minimal media with EA serving as a sole carbon source (NE media). All but an *eutS* mutant exhibited reduced biofilm, where maximum reduction (around 73%) was observed in the case of an *eutN* mutant (Fig. 3). The biofilm in *eutM* and *eutL* mutants were reduced by around 70% and 60%, respectively as compared to WT (Fig. 3). An *eutK* mutant, alternatively produced 51% lower biofilm than WT (Fig. 3).

**Figure 3.**
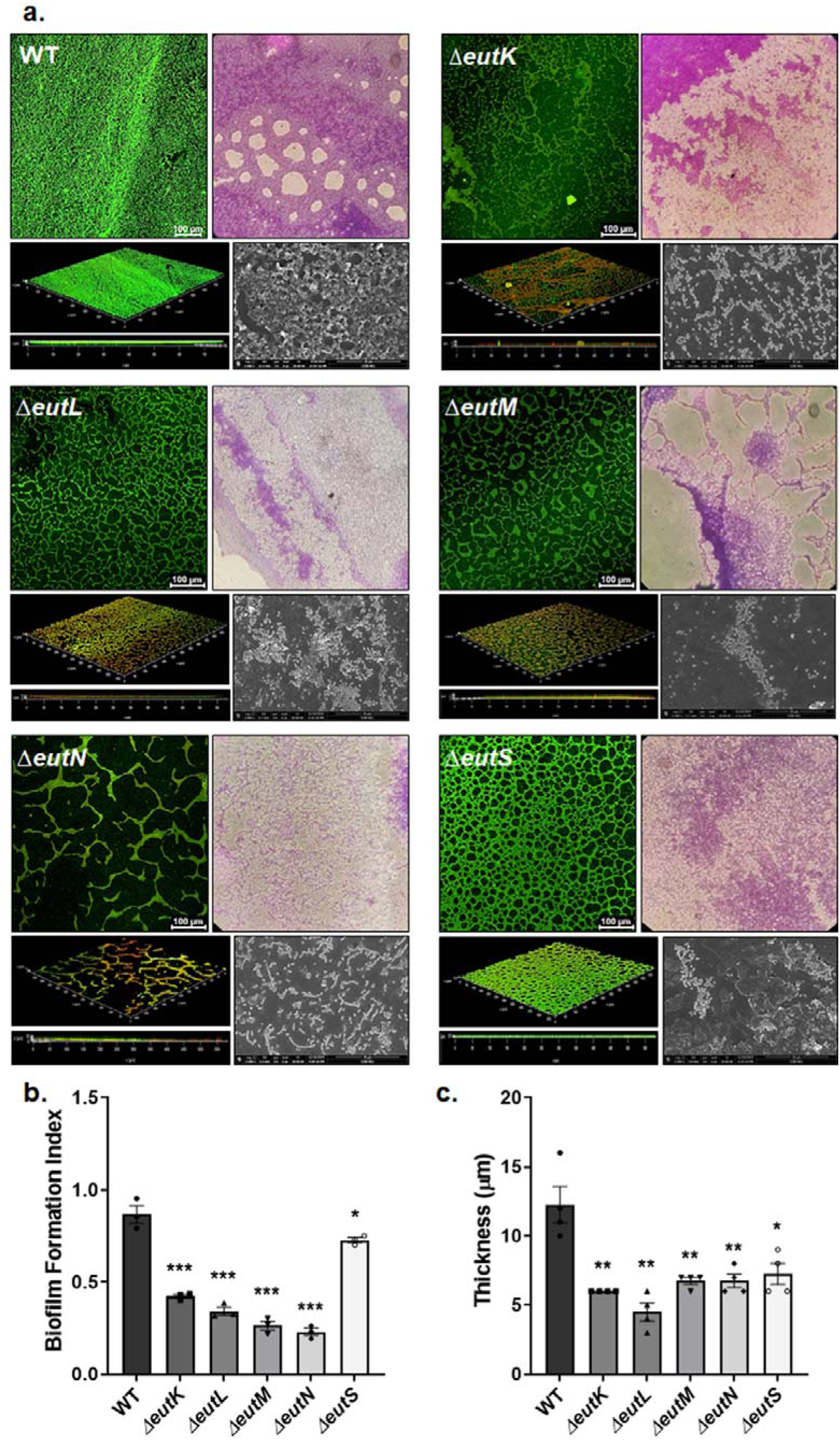
a) Evaluation of biofilm through confocal, crystal violet, and FE-SEM analyses. The samples were stained with SYTO9 (green, live) and propidium iodide (red, dead) for confocal microscopy and visualized under a 20X objective. Orthogonal views of both Z-stacks and 3D images are presented. b) The biofilm formation index (BFI) was calculated using the Crystal Violet assay. c) The thickness of the biofilm was measured in 3D images of the confocal. An unpaired *t*-test (two-tailed) was used for statistical significance. NS p > 0.05, *p < 0.05, **p < 0.01, ***p < 0.001, ****p < 0.0001 compared to the wild type control.

To rule out any polar effect, biofilm was set up with the complemented constructs, which demonstrated a phenotype similar to that of wild-type (Fig. S7a, b), confirming that mutants were responsible for decreased biofilm.

### 3.4 Reduced EA metabolic capacity in the shell protein mutants lowers curli expression in *Salmonella*

Curli expression has a significant impact on biofilm formation. Curli fimbriae and exopolysaccharide (EPS) play crucial roles in cell adhesion and biofilm formation [12, 66–68]. Here, to underpin the effect of the loss of EA metabolism on the morphotype we set out to perform a congo red agar plate assay in rich and minimal media. *Salmonella* produces the red and rough morphotype due to the production of extracellular polysaccharides (EPS) and curli fimbriae [66]. The addition of EA and Vit. B_12_ in TYE and glycerol media produced a dark red colony compared to TYE and glycerol media alone (Fig. 4a, b). In contrast, mutations in the key metabolic enzymes produced a colored colony (Fig. 4a, b), suggesting a strong relationship between EA utilization, curli expression and EPS production. Remarkably, an *eutKM* mutant generating a broken shell produced a light colony (Fig. 4a, b). To investigate the effect of mutations of individual shell proteins, the assay was performed in NE media, where EA served as a carbon source. It was evident that *eutL, eutM,* and *eutK* mutants produced light colonies (Fig. 4c) compared to the wild-type. Out of the three, the *eutL* mutant exhibited strong phenotype (Fig. 4). In contrast, an *eutN* mutant did not grow well in NE media; however, it generated dark red morphotype (Fig. 4). The phenotypes of all individual shell protein mutants were reversed upon complementation with plasmids carrying the respective genes. (Fig. S8)

**Figure 4.**
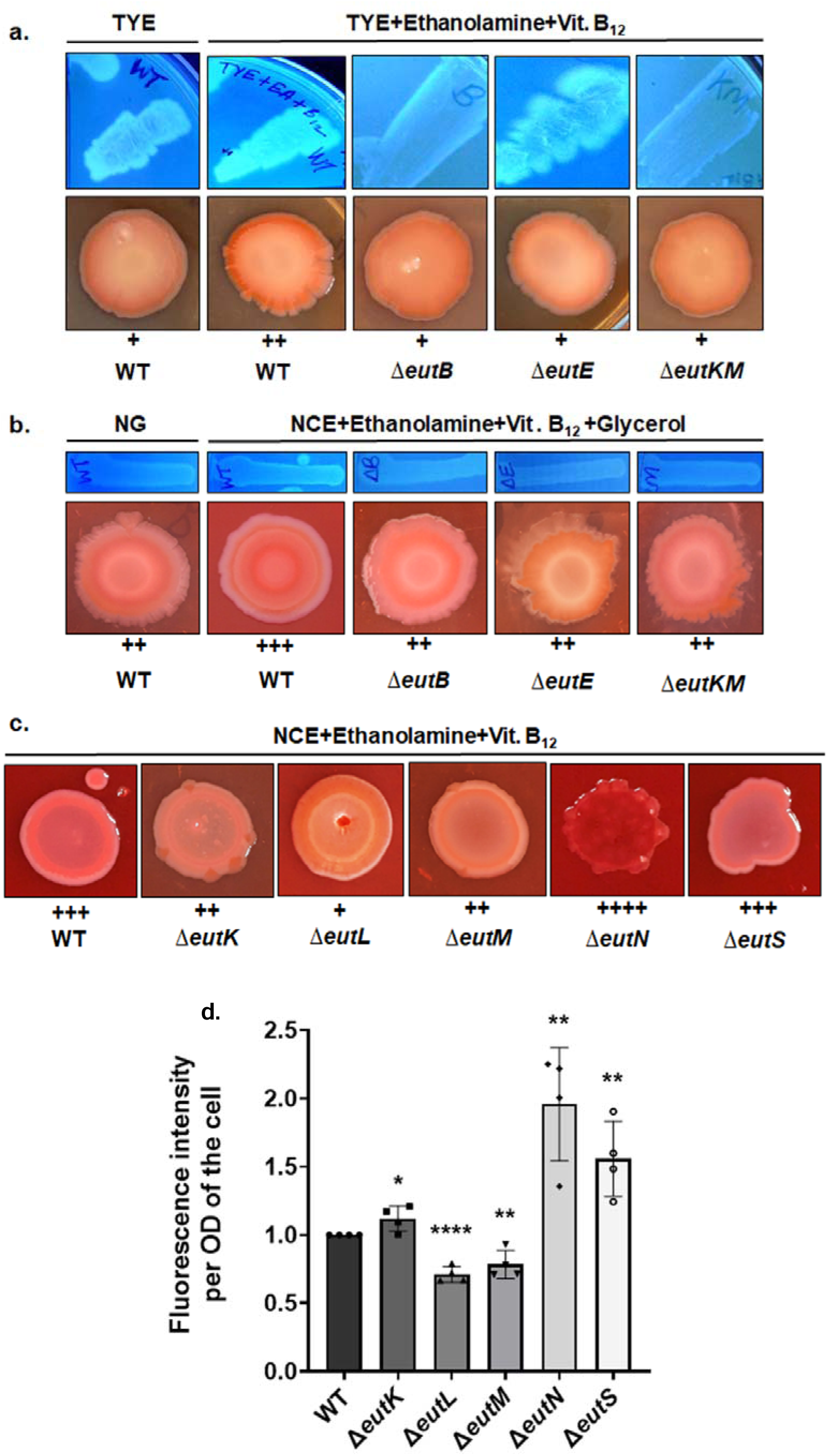
Morphotypes and EPS production of *S.* Typhimurium after 96 h incubation at 37°C on Congo Red agar and Calcofluor Agar plates. Morphotypes and EPS production in WT and the mutants where EA metabolism was completely abolished were evaluated in a) TYE and TYE Supplemented with 30 mM ethanolamine and 150 nM Vit. B_12_ and in b) NCE+Glycerol and NCE+Glycerol supplemented with 30 mM Ethanolamine and 150 nM Vitamin B_12_. c) The morphotypes of WT and the shell protein mutants were assessed in the media where ethanolamine acts as a carbon source, with the media containing NCE, 30 mM ethanolamine, and 150 nM Vit. B_12_. d) Quantification of EPS produced in the WT and the shell protein mutants using Calcofluor white fluorescent assay. An unpaired *t*-test (two-tailed) was used for statistical significance. *p < 0.05, **p < 0.01, ***p < 0.001, ****p < 0.0001 compared to the wild type control. The extent of red color formation in the Congo red assay were denoted with number of + (plus) sign, where one “+” is the light color and four “+” is the dark one.

### 3.5 An *eutL* mutant produces lower EPS in the minimal media

Calcofluor-white generates blue fluorescence after binding with EPS [62]. WT *Salmonella* fluoresced more in UV when EA and Vit. B_12_ were supplemented in rich and minimal media (Fig. 4a, b). In agreement with the morphotype assay results, the *eutB*, *eutE* and *eutKM* mutants also exhibited lower fluorescence intensity, denoting the role of EA metabolism in EPS production (Fig. 4). To ascertain the effect of individual shell protein mutants on EPS production, the mutants were subjected to grow in NE minimal media supplemented with calcofluor and the fluorescence intensity of the bacterial suspension was quantified using a spectrofluorometer after 72 h. The synthesis of EPS was significantly affected in the *eutL* mutant. This mutant produced around 30% lower EPS than WT (Fig. 4d). An *eutM* mutant, on the other hand, generated 9% lower EPS than WT (Fig. 4d). Intriguingly, an *eutN* mutant produced comparatively higher EPS (40% higher) than WT. Knockout of other shell proteins did not affect EPS production (Fig. 4d).

### 3.6 Swarming motility is reduced in the Eut shell protein mutants

The motility is a determining factor in exerting pathogenicity and virulence of bacteria [69]. Earlier studies revealed that strains lacking flagellar motility exhibit poor biofilm-forming ability [70, 71]. To corroborate the role of EA metabolism on swarming motility, the motility diameters were assessed in WT, *eutE*, *eutB* and *eutKM* mutants. It was observed that addition of EA and Vit. B_12_ enhanced the motility of WT *Salmonella*. Albeit, the presence of EA and Vit. B_12_ did not affect the motility of *eutE*, *eutB*, and *eutKM* mutants, where EA metabolism was lost entirely (Fig. 5c, d), affirming the relationship between EA metabolism and motility. Next, we set out to demonstrate whether the deletion of individual shell proteins affects swarming motility of *Salmonella*. Amongst all the shell protein mutants, *eutN*, *eutK*, and *eutM* had severely impaired motility (Fig. 5). These mutants had reduced motility by 3-fold compared to WT. Both *eutL* and *eutS*, on the other hand, produced two-fold lower swarming motility than WT (Fig. 5a, b). As earlier, the loss of swarming motility in the mutants was reversed upon complementing with the respective genes from the plasmid (Fig. S9).

**Figure 5.**
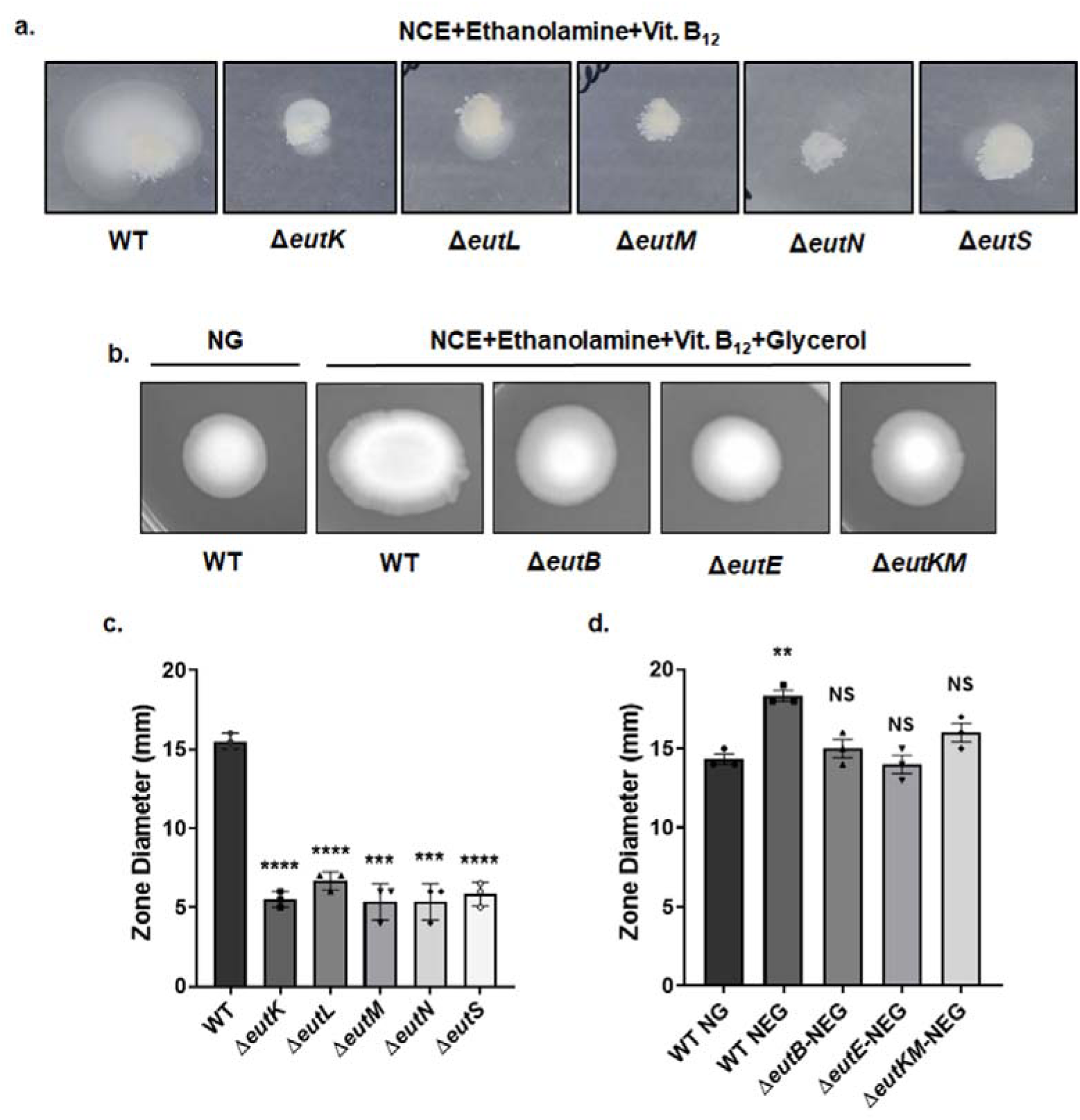
Evaluation of swarming motility in WT and the mutant *Salmonella*. a) and c) The shell protein mutants showed reduced motility as compared to WT when EA was served as carbon source. b) and d) Addition of ethanolamine enhances the motility of WT, whereas the *eutB*, *eutE* and *eutKM* mutant behaved null. An unpaired *t*-test (two-tailed) was used for statistical significance. NS p > 0.05, *p < 0.05, **p < 0.01, ***p < 0.001, ****p < 0.0001.

### 3.7 Genes involved in biofilm, motility, and invasion are downregulated in the shell protein mutants

To establish the relationship of EA metabolism with genes related to biofilm, motility and invasion, we sought to determine the expression of *csgD*, *fljB* and *sipA* in minimal media. It was observed that WT *Salmonella* grown in a minimal medium containing EA and B_12_ had slightly higher expression of all the aforementioned genes than those grown in media without EA as a carbon source (Fig. S10). However, there were no significant changes in the expression of the above genes in the *eutB*, *eutE,* and *eutKM* mutants devoid of enzyme and MCP shell (Fig. S10)

To evaluate if lower biofilm formation in the shell protein mutants was due to the change in the expression of *luxS* and *csgD*, the genes responsible for quorum sensing and curli fimbriae synthesis; the expression profile of the same was determined through qRT-PCR analyses. It was revealed that the maximum downregulations in the *luxS* and *csgD* gene expression were observed in the sessile cells of the *eutL* and *eutM* mutants. The expression of *luxS* was downregulated by 92% and 63% in *eutL* and *eutM*, respectively (Fig. 6). In case of *csgD*, the fold reduction in the respective mutants was 69% and 95% (Fig. 6). By contrast, the *luxS* expression in planktonic cells of *eutL* was reduced by 93% and that in case of *eutM* was 86%.

**Figure 6.**
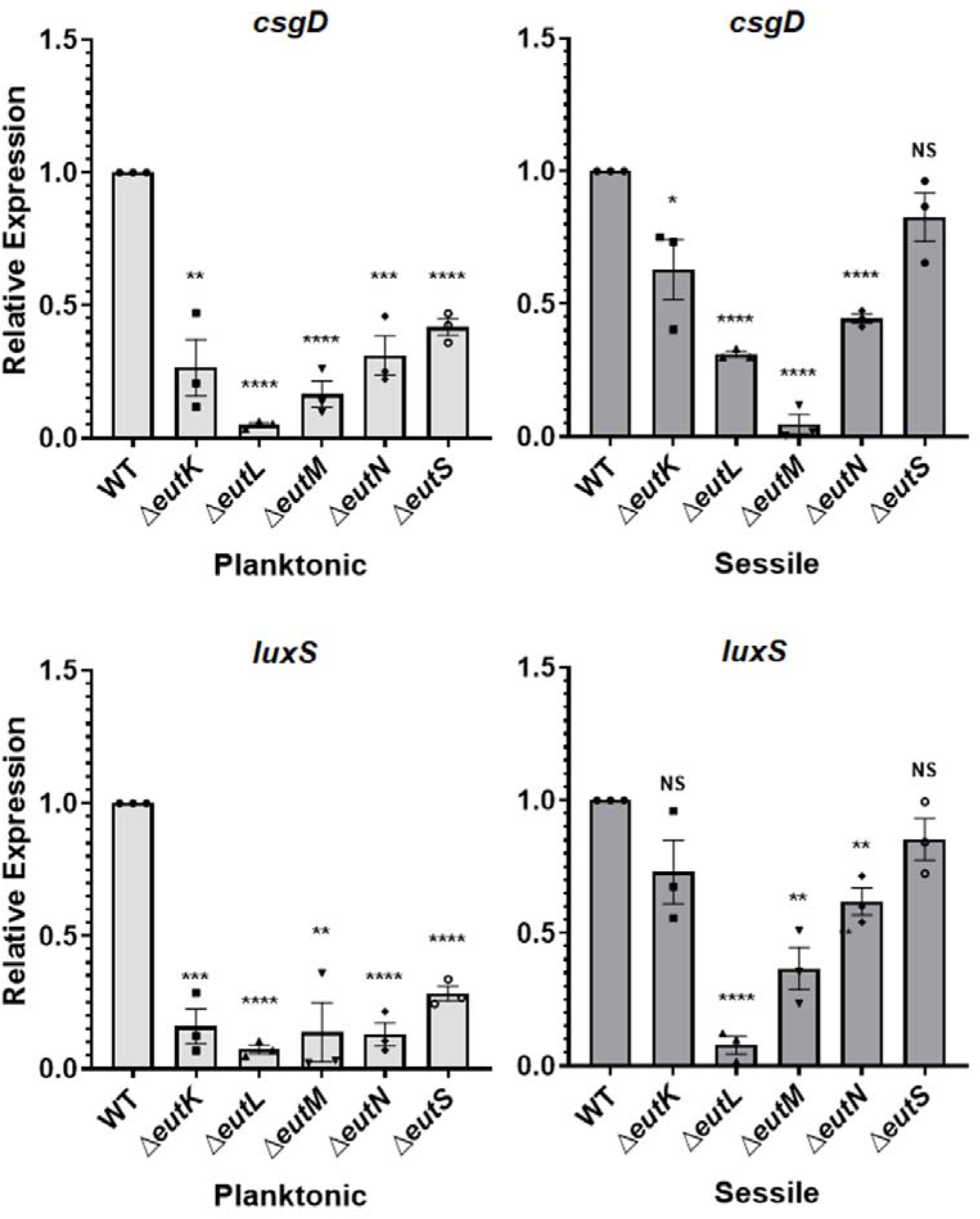
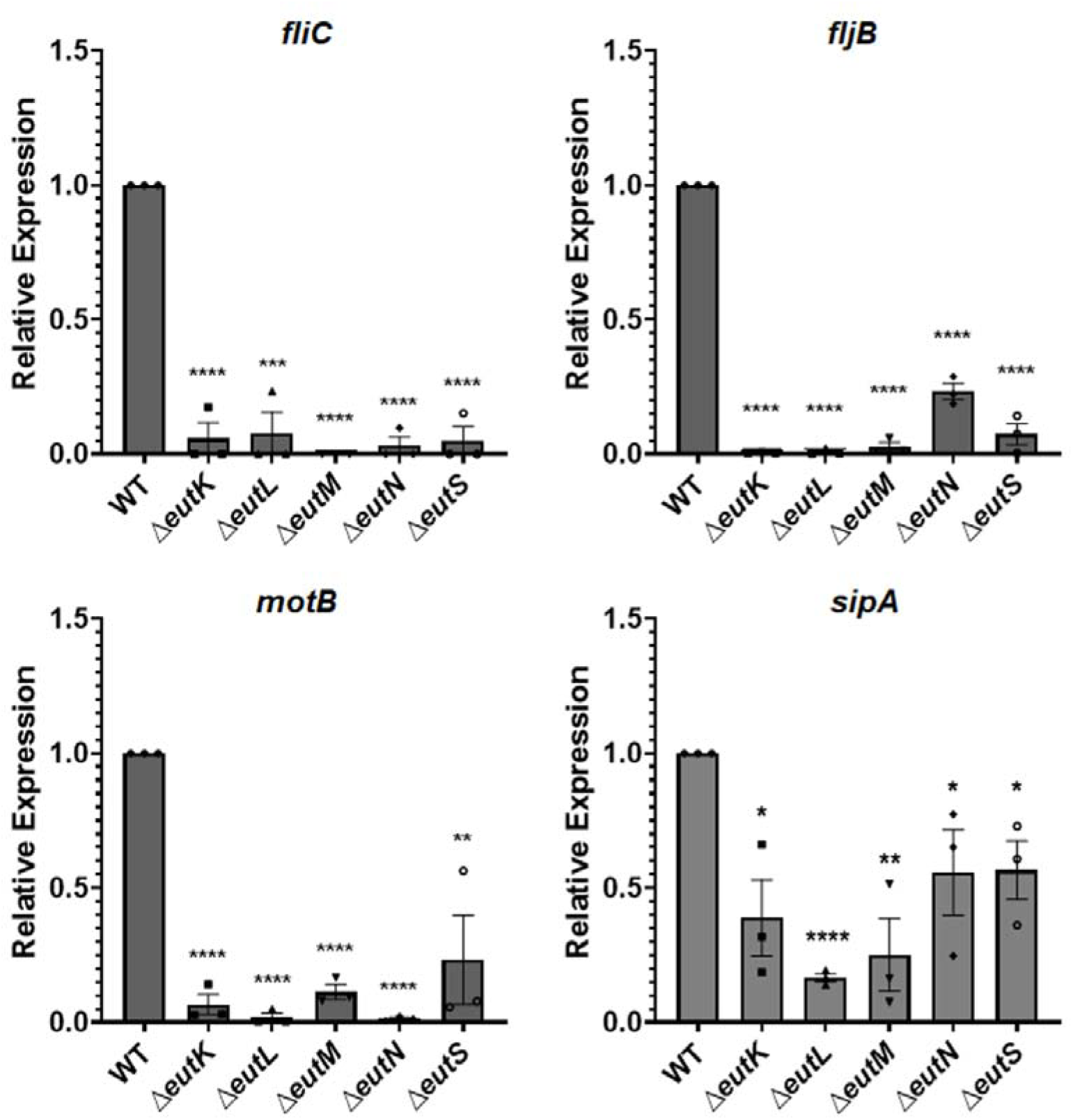
Gene expression analysis of gene targets related to biofilm, motility, and cell invasion. An experiment was performed with three technical replicates and three biological replicates. The pathogenicity related genes were down regulated in almost all the shell protein mutants. An unpaired *t*-test (two-tailed) was used for statistical significance. NS p > 0.05, *p < 0.05, **p < 0.01, ***p < 0.001, ****p < 0.0001.

At the same time, *csgD* expressions in the planktonic *eutL* and *eutM* cells were downregulated by 95% and 83%, respectively (Fig. 6).

We also set out to determine the expression of motility and invasion-related genes. It was indicated that the expression of *fliC*, *fljB*, and *motB* was downregulated by more than 80% in all the shell protein mutants (Fig. 6). Whereas, maximum downregulation (83%) of *sipA* expression was observed in the case of an *eutL* mutant.

### 3.8 An *eutL* mutant is less viable than WT in the intramacrophagic environment

Since expression study indicated maximum downregulation of the *sipA* gene in an *eutL* mutant, we sought to examine the intramacrophagic survival potential of the *eutL* mutant. As the ethanolamine utilization operon is known to be inhibited by glucose (64), a modified no-glucose DMEM was used, supplemented with 1 mM pyruvate during infection. In the current study, invasive defective *Salmonella* [21, 22] were used by constructing an *invA* mutation in the background of WT and mutant strains. To visualize bacteria inside the macrophage RAW264.7, the wild-type *Salmonella* and mutants were transformed with a plasmid expressing m-Cherry (pLac22-mCherry). A red fluorescent protein, m-Cherry, was expressed after adding 1 mM IPTG. The bacteria within the macrophages were visualized under a fluorescent microscope. To determine the counts of phagocytosed bacteria, the macrophages were washed and subjected to lysis with 0.1 % Triton X-100. Serial dilutions were plated to determine the viable bacteria as colony-forming units (cfu)/ mL of the cell lysates. Here, viable CFUs at the indicated time points were defined as the percentage of this intracellular time 0 h population and normalized in such a way that the wild-type was equal to 100%, as depicted earlier [21]. It has been observed that the number of recovered bacteria was reduced to 50% in the case of the *eutL* mutant (Fig. 7a, b). An *eutE* mutant, defective in producing acetaldehyde dehydrogenase, had only 40% viable cells remaining (Fig. 7a, b)

**Figure 7.**
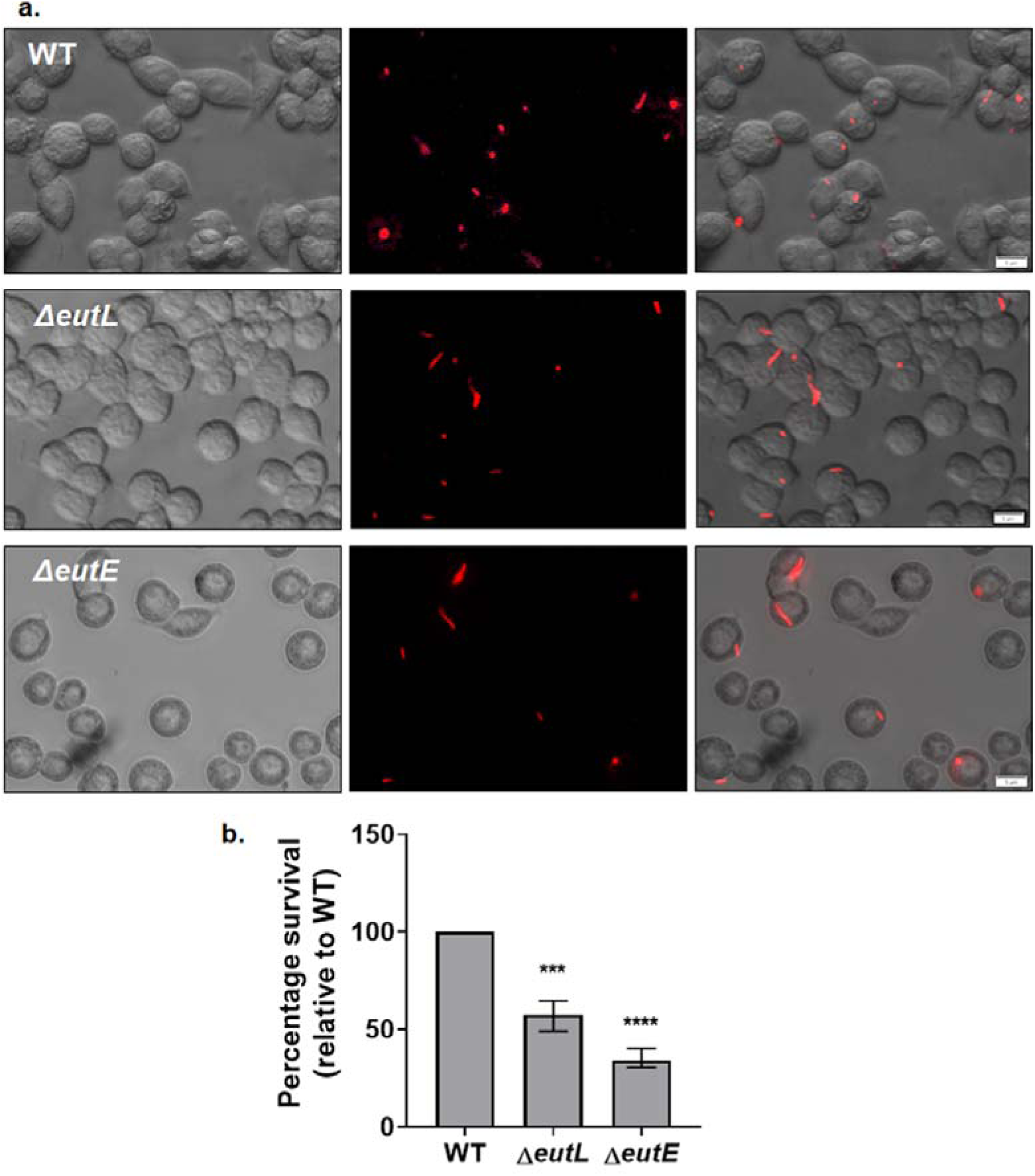
Visualization of intracellular bacteria within macrophage cells RAW264.7. a) Infection of the RAW264.7 cells with WT, *eutL*, and *eutE* mutant strains of *Salmonella* (50:1 MOI) expressing m-Cherry (red). A representative microscopy merged image shows a bright field with bacteria expressing m-Cherry. The imaging was done at four hours post-infection. b) Percentage of recovered bacterial cells after four hours of infection. The experiments were done in triplicate with three biological replicates. An unpaired *t*-test (two-tailed) was used for statistical significance. *p < 0.05, **p < 0.01, ***p < 0.001, ****p < 0.000.

## 4. DISCUSSION

Despite several advances in chemotherapy, non-typhoidal *Salmonella* (NTS) has become one of the significant threats to global public health due to the prevalence of multi-drug resistant (MDR) variants. In a list of global priority pathogens for antimicrobial research published by the World Health Organization (WHO), fluoroquinolone and third-generation cephalosporine-resistant variants have rendered major cause for concern (https://www.who.int/news/item/27-02-2017-who-publishes-list-of-bacteria-for-which-new-antibiotics-are-urgently-needed). Additionally, intracellular *Salmonella* Typhimurium has been found resilient to aminoglycoside antibiotics [72]. Despite numerous antimicrobial compounds discovered so far to mitigate the pathogen, only few cellular processes are targeted to design new drugs [73]. It is suggested that Eut microcompartment (MCP) metabolism could be a potential template for designing novel drugs in order to combat multidrug resistant (MDR) pathogens [30, 31]. Eut MCP metabolism is an essential genomic determinant of pathogenicity associated with gastroenteritis [28], and factors affecting Eut MCP metabolism might interfere with bacterial pathogenesis [31]. Although several advances in understanding EA metabolism in *Salmonella* have been made so far (26, 27, 42, 46, 49, 51), the lack of understanding the significance of EA metabolism in bacterial physiology limits the progress in developing novel therapeutics affecting microcompartment metabolism.

The apparent role of EA metabolism in modulating bacterial physiology, viability and intrinsic antibiotic resistance could not be established in the nutrient rich media, the condition where most modern therapeutics are developed [74]. Interestingly, the phenotype was profoundly manifested under minimal media conditions, which created a paradigm shift underscoring the role of Eut MCP metabolism in modulating bacterial physiology. Minimal media supplemented with EA provides a validated approximation of the gut environment in which *Salmonella* disseminates during inflammation and exemplifies the roles of specific nutrients in the bacterial physiology. The use of minimal media to explore novel antimicrobial targets was also depicted elsewhere [18, 20, 64, 74].

Since the mutation of individual shell protein exhibited small phenotype in the conventional MIC determination, we conducted growth analysis to capture minor fitness effects of mutations of all the individual shell proteins in the presence of structurally unrelated antibiotics [64, 65, 75]. The antibiotic susceptibility index of the mutants relative to wild-type (ASI) indicates that the deletion of *eutL* and *eutM* impart a significant impact on the intrinsic antibiotic resistance of *Salmonella*, where maximum sensitivity is observed in an *eutL* mutant (Fig. 2). It is tempting to observe that genetic mutations on the *eutL* and *eutM* result the bacterial cell susceptible mostly to ciprofloxacin, chloramphenicol and kanamycin - the antibiotics that interfere DNA replication and translation (Fig. 2). In *Salmonella*, EA serves as a source of carbon, nitrogen and energy. A disruption in EA metabolism in *Salmonella* not only impairs the several important biosynthetic pathways but also hampers ATP generation; which might exert more impact on nucleic acid metabolism.

EutL and EutM are structurally orthologous to PduB and PduA shell proteins of 1, 2-propanediol utilization microcompartment (Pdu MCP) [25, 47]. The former forms a pseudohexameric trimer and a canonical hexamer [25, 47]. Eut MCPs of *Salmonella* had not been purified yet. However, previous literature suggested that hexameric and trimeric shell proteins share the maximum percentage of total shell protein content (68), where trimer alone contributes around 50% amongst all shell proteins [76]. The trimeric EutL was crystallized in both open and closed conformations and was proposed to allow the transport of large molecules through its central pore [47, 52]. EutL also interacts with ethanolamine substrate [52]. Alternatively, the hexameric EutM shell protein is presumed to transport the EA substrate [77]. A recent study revealed that individual deletion of either EutM or EutL shell protein results in a growth defect and formation of polar aggregates [51]. We presumed that EA metabolism might have been disrupted in *eutL* and *eutM* mutants due to aberrant MCP shell assembly. The release of high level acetaldehyde intermediate (Fig. S5) in the *eutL* and *eutM* mutant indicates severe carbon loss, which supports our hypothesis.

It is also revealed that biofilm formation, curli expression and EPS production are reduced in the cell where either of the major MCP shell proteins are lost (Fig. 3). We surmise that EA metabolism upregulates pathogenicity related genes present in *Salmonella* pathogenicity island-1 (SPI-1) that enhances physiological fitness (Fig. S6). It can be presumed that the diminished biofilm and EPS in the shell protein mutants increase the permeability of antibiotics, thereby lowering the intrinsic antibiotic resistance in those mutants. Recently, genetic disruption of MCP formation has been shown to repress Fusobacterial pathogenesis and virulence [78]. Surprisingly, bacterial cells devoid of EutN shell protein was observed to gain resistance to some antibiotics as determined from the ASI profile (Fig. S2). The heightened EPS in the *eutN* mutant (Fig. 4) might presumably inhibit the entry of antibiotics, resulting transient resistant phenotype. Nevertheless, an *eutN* mutant exhibits severe biofilm phenotype (Fig. 3), presumably due to poor growth of the said mutant [51, 78]. It is worth noting that the current study with *Salmonella* Typhimurium demonstrates the relationship between EA metabolism and biofilm, when EA is employed as carbon source. An in-depth analysis of preferential nutrients indicated that *Salmonella* produces more biofilms when EA is consumed as sole carbon source (Fig. S11); unlike adherent-invasive *Escherichia coli*, which prefers EA as nitrogen source [36]. Likewise, phenotype determining the effect of individual shell protein mutations on physiology and intrinsic antibiotic resistance of *Salmonella* is greatly manifested when EA is lone source of carbon.

Further, inactivation of EA metabolism in one of the major shell protein mutants also renders *Salmonella* susceptible to the host’s innate defense system (Fig. 7), suggesting the relationship between EA metabolism and intramacrophagic survival. The role of MCP-mediated EA utilization in the intracellular survival and replication of *Listeria monocytogenes* has been demonstrated recently [79].

As a whole, genetic mutation targeting major shell proteins disrupting EA metabolism would thus provide significant resources in designing future antimicrobials against *Salmonella*. To further validate the strategy to an antibiotic resistant variant, we set out to examine the effect of deletion of a major shell protein, EutL, on the growth of a chloramphenicol-resistant *Salmonella*. In this study a *cam* cassette was inserted at the *invA* chromosomal locus (*invA*::*frt-cam-frt*) of both WT and an *eutL* mutant. The bacteria were grown in minimal media supplemented with EA and B_12_, and the growth was monitored in the microplate reader, as mentioned in the methodology section. It was observed that the OD_600_ of the cell containing secondary mutation at *eutL* declined at a comparatively faster rate at increasing chloramphenicol concentration than the cell having EutL shell protein intact (Fig. S12), indicating higher antibiotic susceptibility of the *eutL* mutant.

Furthermore, the current study also provides deep insights into understanding the differential roles of shell proteins in Eut MCP assembly and metabolism.

## 5. CONCLUSION

In this seminal work multifaceted roles of Eut MCP in supporting the pathogen’s physiology and virulence are elucidated. Genetic disruption of major shell proteins of Eut microcompartment not only abrogates ethanolamine metabolism in *Salmonella* but also hinders biofilm and curli expression. The reduced physiological fitness in the shell protein mutants lowers intrinsic resistance of *Salmonella* to some antibiotics and reduces survival against host immune responses. This observation demonstrates that the integrity of the Eut MCP plays a critical role in bacterial pathogenicity and could be exploited as a potential therapeutic target. The further research is required to evaluate the underlying role of MCP integrity on ethanolamine metabolism of *Salmonella* through *in vivo* animal model system to design treatment regimens.

## SUPPLEMENTAL MATERIAL

Supplementary data associated with this article can be found in the online version.

## Supporting information

Supplementary data associated with this article can be found in the online version

## ACKNOWLEDGMENTS

This study was supported mainly by the ad-hoc grant from the Indian Council of Medical Research (ICMR) [Diarr/Adhoc/5/2022-ECD-II] to C.C. and A.S.G. The Authors also thank the SERB-Ramanujan fellowship grant (SB/S2/RJN-128/2017) for initial support. M.R.B. and A.S.D. acknowledge the Council of Scientific and Industrial Research (CSIR) and the Department of Biotechnology, Govt of India (DBT), respectively, for their fellowships. The authors thank Ms. Yashodhara Shinde for her tireless help in constructing secondary mutations at the *invA* gene in WT, *eutL*, and *eutE* mutant cells. Authors also thank Mr. Rohit Dashpute for making an *eutB* mutant construct. The authors thank Dr. Syed G. Dastager and Dr. Mahesh S. Dharne of National Collection of Industrial Microorganisms (NCIM), CSIR-National Chemical Laboratory, Pune, India, for their help in microplate reader access, qRT-PCR, nucleotide sequencing and other services.

## AUTHOR CONTRIBUTIONS

**MRB**: Performed bacterial genetics, physiology and molecular biology work, carried out microscopic visualization and data analyses, wrote original draft; **ASD**: Performed molecular biology work, gene expression analysis, confocal microscopic analyses; **ARS**: Cell culture work and fluorescence microscopy; **ASG**: Investigation, formal analysis of cell culture work and funding acquisition; **TNH**: Performed molecular biology work; **TAB**: Supervision, formal analyses and initial resources; **AS**: Biofilm work; **SBA**: Supervision, **CC**: Conceptualization, initial molecular biology and genetics work, Supervision, Funding acquisition, Project administration, Supply of resources, Writing - review & editing.

## DECLARATION OF COMPETING INTERESTS

The authors declare no conflict of interest.

## Notes

### Competing Interest Statement

The authors have declared no competing interest.

### Summary of Updates

Middle name of author (Dr. Sachin B. Agawane) has been corrected. Figure 4. has been edited

