## Supplementary data associated with this article can be found in the online version for "Deletion of major shell proteins of ethanolamine utilization microcompartment reduces intrinsic antibiotic resistance, biofilm, and intracellular survival of *Salmonella* Typhimurium"

**Tables**

**Table S1**. Bacterial Strains used in the study

| **Species/Strain** | **Genotype** | **Source** |
| --- | --- | --- |
| ***E. coli*** | | |
| **YS9** | DH5alpha/pKD46 | Lab Collection |
| **YS10** | DH5alpha/pCP20 |  |
| **MB1** | DH5alpha/pLac22 |  |
| **CE28** | *E. coli*/ pKD4 |  |
| **CE29** | *E. coli* BW25141/pKD3 |  |
| ***Salmonella enterica* Typhimurium LT2** | | |
| **CE1** | LT2 *Salmonella enterica* serovar Typhimurium LT2 | Lab Collection |
| **CE35** | ∆*eutK*::frt |  |
| **CE34** | ∆*eutL*::frt |  |
| **CE37** | ∆*eutM*::frt |  |
| **CE36** | ∆*eutS*::frt |  |
| **CE3** | LT2/pLac22 |  |
| **RD8** | ∆*eutB*::frt | This Study |
| **MB7** | ∆*eutN*::frt |  |
| **AD101** | ∆*eutE*::frt |  |
| **AD43** | ∆*eutK*::frt-∆*eutM*::frt-Cam-frt |  |
| **AD24** | Δ*eutK*::frt/pLac22 |  |
| **AD8** | ∆*eutK*::frt/pLac22-*eutK* |  |
| **MB34** | Δ*eutL*::frt/pLac22 |  |
| **MB33** | ∆*eutL*::frt/pLac22-*eutL* |  |
| **AD25** | Δ*eutM*::frt/pLac22 |  |
| **AD26** | Δ*eutM*::frt/pLac22-*eutM* |  |
| **MB36** | Δ*eutN*::frt/pLac22 |  |
| **MB30** | Δ*eutN*/pLac22-*eutN* |  |
| **MB38** | Δ*eutS*::frt/pLac22 |  |
| **MB37** | Δ*eutS*/pLac22-*eutS* |  |
| **YS46** | Δ*invA*::*frt-cam-frt* |  |
| **YS47** | Δ*invA*::*frt* |  |
| **YS56** | Δ*eutL::frt-*Δ*invA*::*frt-cam-frt* |  |
| **YS61** | Δ*eutE::frt-*Δ*invA*::*frt-cam-frt* |  |

**Table S2**. Primers used in this study

| **Description** | **Primer** | **Sequence (5’-3’)** |
| --- | --- | --- |
| **Mutant Preparation** | | |
| ***eutK*** | *eutK-Cam-F* | TGCTGGATATCGCCCGTAACCCTGTCCAGCGTGCGTAACGGAGGCTGCCATGTAGGCTGGAGCTGCTTCG |
|  | *eutK-Cam-R* | CATGATCTTCTATGCCGGACGCCCGCCAGGACGCCCGGCCCTCCGACAGAATATGAATATCCTCCTTAGTTC |
| ***eutL*** | *eutL-Cam-F* | TGCTGGAGCAGAAAGCGTCCGGCATCAACATGACCCGTTAAGGAGACATCTGTAGGCTGGAGCTGCTTCG |
|  | *eutL-Cam-R* | ATTCCGTCCACTTCCAGTAATCCCAGGGCATTGATCATTGGCAGCCTCCGATATGAATATCCTCCTTAGTTC |
| ***eutM*** | *eutM-Cam-F* | GGTTGAATGAACGGTCCCGTTCTGGACCCCTTTAATGAGAGGAAAACACGTGTAGGCTGGAGCTGCTTCG |
|  | *eutM-Cam-R* | GCGCGCATTCATGCCGGATGGCGACGCTGCGCGTCTTATCCGGCCTACCAATATGAATATCCTCCTTAGTTC |
| ***eutN*** | *eutN-Cam-F* | CATTGCCTGATGGTGCGCAGATTCAAGGCCTACGACCCAACGGGCAATCCTGTAGGCTGGAGCTGCTTCG |
|  | *eutN-Cam-R* | TACCGCTTTCACCACCTGTTCAATATCCTGTTGATTCATGATGTTCAATCATATGAATATCCTCCTTAGTTC |
| ***eutS*** | *eutS-Cam-F* | CCGACAAAAAATTGCCACGATGACGGCAGTTTCAGTGGAGACGGTGAGCATGTAGGCTGGAGCTGCTTCG |
|  | *eutS-Cam-R* | CCGGCACCGACCGCGCCGACAAAAGCAATACGTTTCATTACCGCCACCAAATATGAATATCCTCCTTAGTTC |
| ***eutB*** | *eutB-Cam-F* | GGTGAAATCACTCGCATTTCCTTCCTGAGGGAACGACTTTGTAGGCTGGAGCTGCTTCG |
|  | *eutB-Cam-R* | TCAATCTGTTTTTGATCCATGGTGTTATCCCCGCGTCAATATGAATATCCTCCTTAGTTC |
| ***eutE*** | *eutE-Cam-F* | CTGGCGGGAAAGTGGTTTTCCATAAATAGGATTGAACATCTGTAGGCTGGAGCTGCTTCG |
|  | *eutE-Cam-R* | CGTGCGCCATGAGTCATCCCTTATACAATGCGAAACGCATATATGAATATCCTCCTTAGTTC |
| ***invA*** | *InvA-Cam-F* | TGAAAAGCTGTCTTAATTTAATATTAACAGGATACCTATA TGTAGGCTGGAGCTGCTTCG |
|  | *InvA-Cam-R* | GTTTTTATAACATTCACTGACTTGCTATCTGCTATCTCAC ATATGAATATCCTCCTTAGTTC |
| **Mutant and clone confirmation** | | |
| **pLac** | PLac-F | CCCCAGGCTTTACACTTT |
|  | PLac-R | GTTAGATTTCATACACGGTGCCTG |
| ***eutK*** | *eutK*-F | TCGTACCTCTCGTCTACCGCA |
|  | *eutK*-R | TCATCAAGCAGGATTTCCGT |
| ***eutL*** | *eutL*-F | GGCAGTCGGAAAGCCTTT |
|  | *eutL*-R | CGACTAATCACACGGCCTGT |
| ***eutM*** | *eutM*-F | AAATCACTCAGCGTCTGGGA |
|  | *eutM*-R | TCATTTCCACCATCAGCAAT |
| ***eutN*** | *eutN*-F | TGCAAAGCCGCAACCGAT |
|  | *eutN*-R | TTTCGCTGACGGCAAGTT |
| ***eutS*** | *eutS*-F | CGGAACAGGCTTAAATTCAGA |
|  | *eutS*-R | TGCAACGTGGTAATTAAGGC |
| ***eutB*** | *eutB -F* | TACGCCTCTTTTTGGCGGATCGGTT |
|  | *eutB -R* | CTGTCCCATTGACGCCATCACGCTA |
| ***eutE*** | *eutE-F* | ATCGACAGTATCGGGGCGGG |
|  | *eutE-R* | ACCACCATCGACACCACGTC |
| **qRT PCR** | | |
| Gyrase  (Reference) | *gyrA-F* | CTCTACTCCCAGACCCAGCT |
|  | *gyrA-R* | GCTTCAAGGATATGCGCACG |
| Transcriptional regulator of curli fimbriae synthesis | *csgD-F* | GCCATAACCGGAAAACTGCA |
|  | *csgD-R* | CCACGTGTTCCTGGTCTTCA |
| Flagellar Filament Flagellin | *fliC-F* | ACACTGCAGATGACGGTACA |
|  | *fliC-R* | CGGGTTTTCGGTGGTTGTAG |
|  | *fljB-F* | ATCCAGGCTGAAATCACCCA |
|  | *fljB-R* | GTTCAGTGAGTCCAGACCCA |
| Bacterial Quorum Sensing | *luxS-F* | GAACAAAGAAGTGATGCCGGA |
|  | *luxS-R* | TAAACGTTCAGCTCCGGGAT |
| Flagellar Motor Rotation | *motB-F* | GCCTACGCCGATTTTATGAC |
|  | *motB-R* | CTGTTGGGTGTAATCATCGC |
| Cell Invasion Protein | *sipA-F* | AGCGTGGATAACAGTAAGCA |
|  | *sipA-R* | CACACGCGAATGACTATGAC |

| **Antibiotic (µg/mL)** | **WT** | **Δ*eutK*** | **Δ*eutL*** | **Δ*eutM*** | **Δ*eutN*** | **Δ*eutS*** |
| --- | --- | --- | --- | --- | --- | --- |
| Ciprofloxacin | 0.016 | 0.016 | 0.012 | 0.023 | 0.016 | 0.016 |
| Cefotaxime | 0.094 | 0.094 | 0.094 | 0.094 | 0.094 | 0.094 |
| Cephalothin | 3 | 3 | 3 | 3 | 1.5 | 1.5 |
| Chloramphenicol | 3 | 3 | 4 | 2 | 2 | 3 |
| Piperacillin | 1.5 | 2 | 2 | 2 | 1.5 | 2 |
| Kanamycin | 32 | 32 | 24 | 24 | 32 | 32 |
| Ampicillin | 1.5 | 1.5 | 2 | 1.5 | 1.5 | 1.5 |

**Table S3a:** Minimum inhibitory concentrations (MIC) of various unrelated classes of antibiotics as determined by Ezy MIC strip (Himedia) method.

**Table S3b:** Minimum inhibitory concentrations (MIC) of various unrelated classes of antibiotics as determined by broth dilution method

| **Antibiotic (µg)** | **WT** | **Δ*eutK*** | **Δ*eutL*** | **Δ*eutM*** | **Δ*eutN*** | **Δ*eutS*** |
| --- | --- | --- | --- | --- | --- | --- |
| Ciprofloxacin | 0.027 | 0.013 | 0.013 | 0.013 | 0.013 | 0.013 |
| Cefotaxime | 0.156 | 0.156 | 0.156 | 0.156 | 0.156 | 0.156 |
| Cephalothin | 2.5 | 2.5 | 2.5 | 2.5 | 2.5 | 2.5 |
| Chloramphenicol | 2.5 | 2.5 | 2.5 | 2.5 | 2.5 | 2.5 |
| Piperacillin | 0.625 | 0.625 | 0.625 | 0.625 | 0.625 | 0.625 |
| Kanamycin | 50 | 50 | 50 | 50 | 50 | 50 |
| Ampicillin | 0.375 | 0.375 | 0.375 | 0.375 | 0.375 | 0.375 |

*Experiments in broth dilution method were carried out with four technical replicates with three biological replicates, and the most reproduced results have been shown.

**FIGURES**

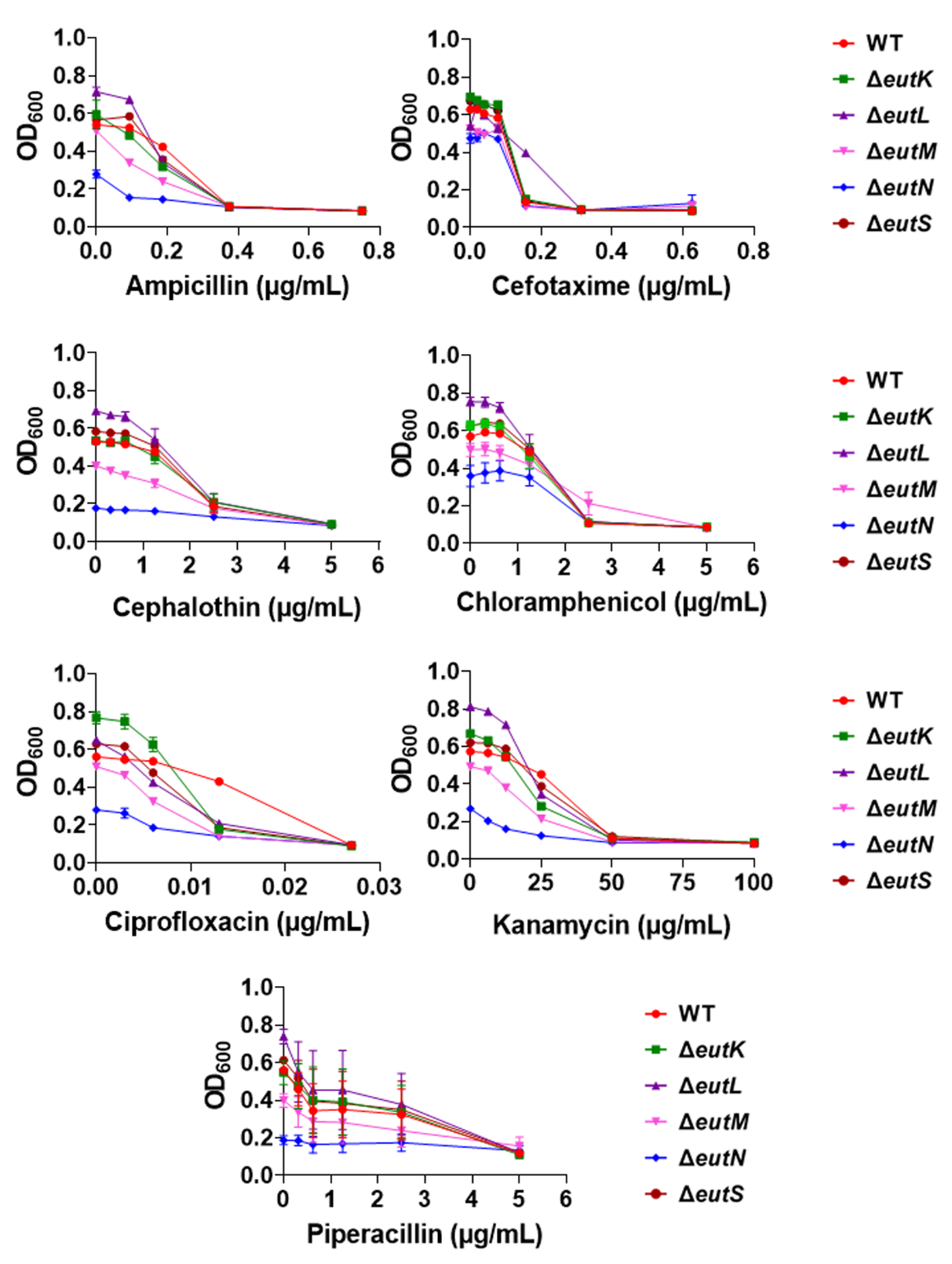

**Figure S1**. The effect of the various antibiotics listed was evaluated against WT and shell protein mutants as defined by growth measured by optical density (OD_600_) after 24 h at 37°C.

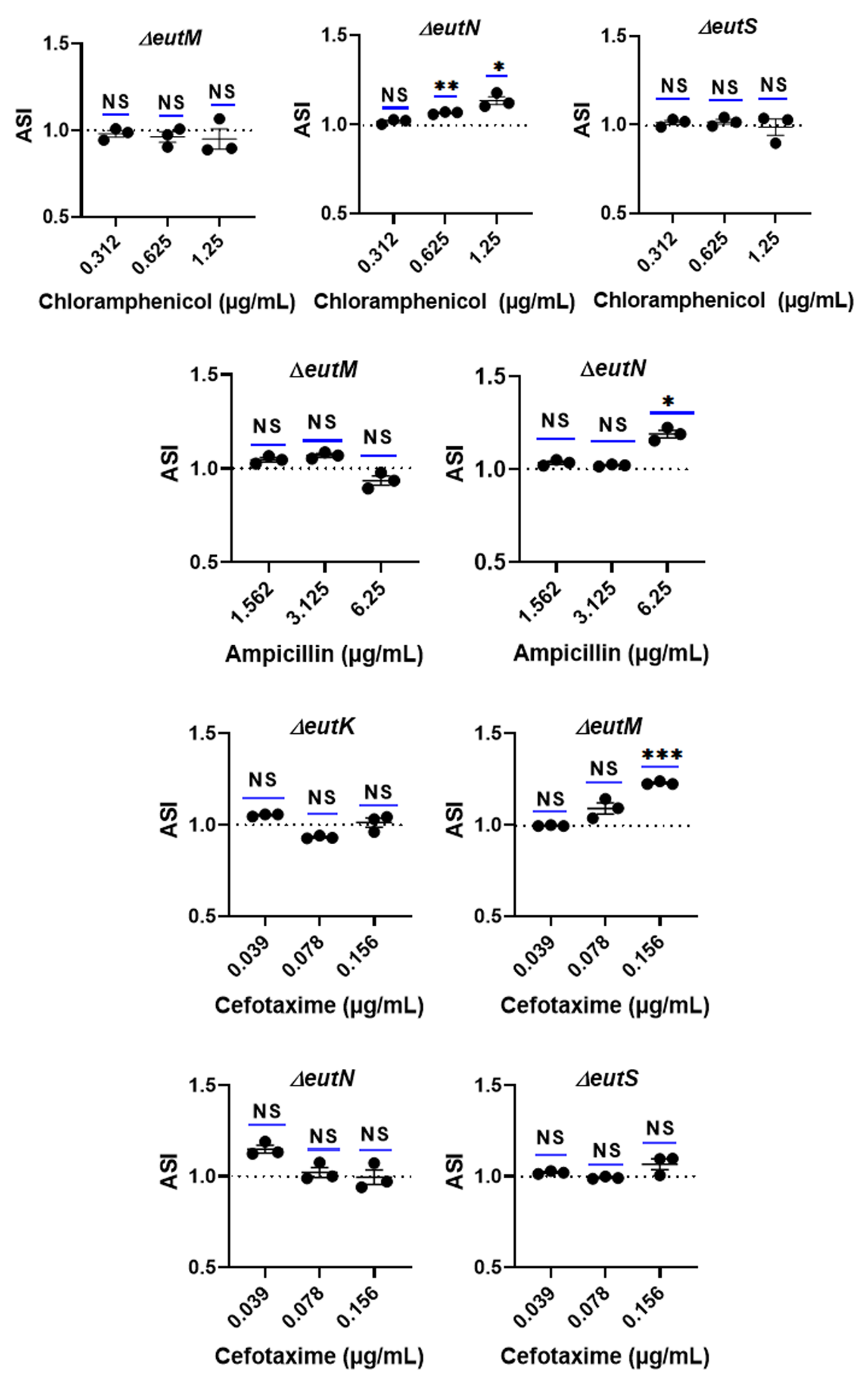

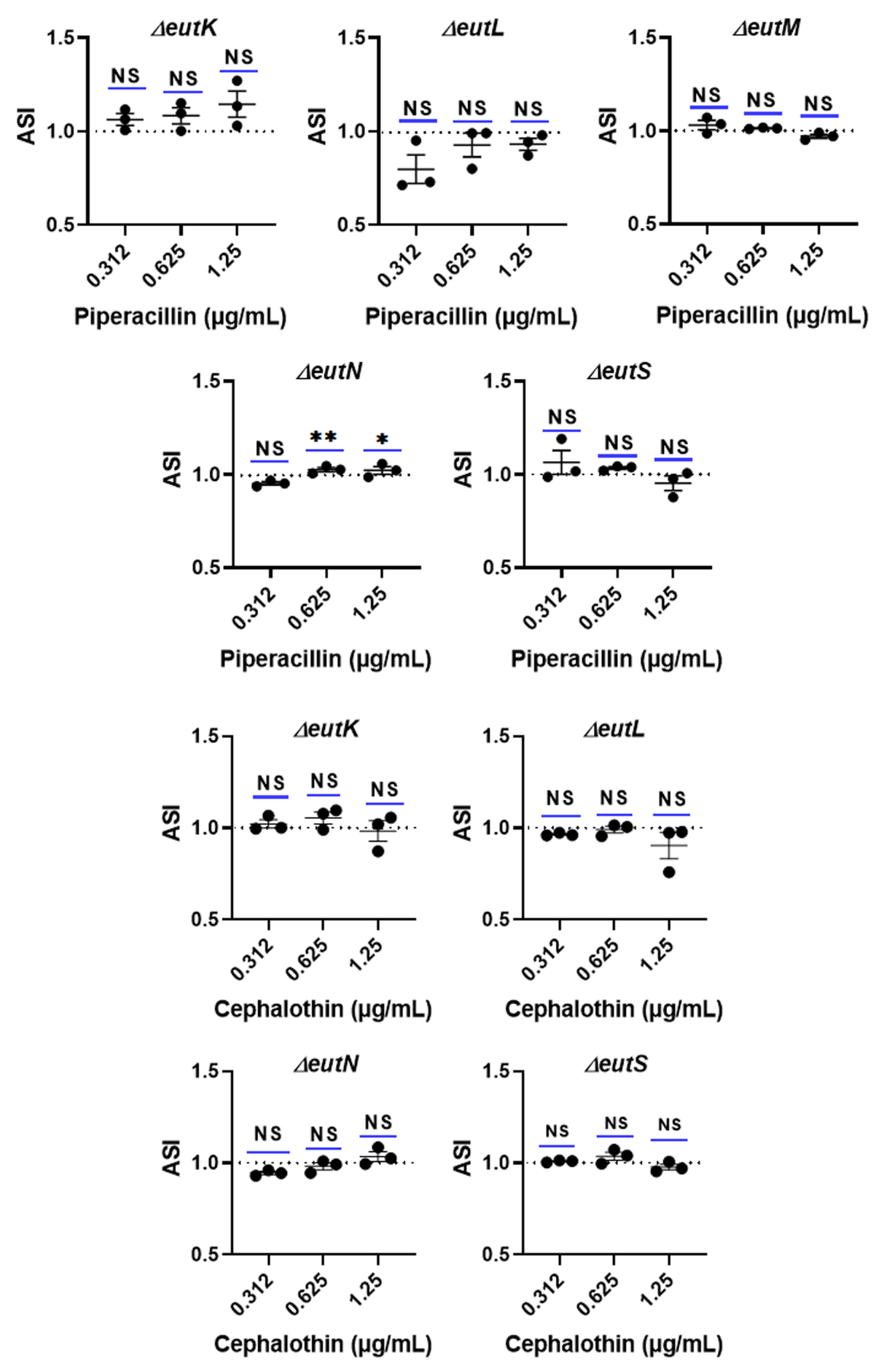

**Figure S2.** Antibiotic susceptibility Index (ASI) of shell protein mutants in presence of different antibiotics. Mutants, which were not susceptible to antibiotics are shown here.

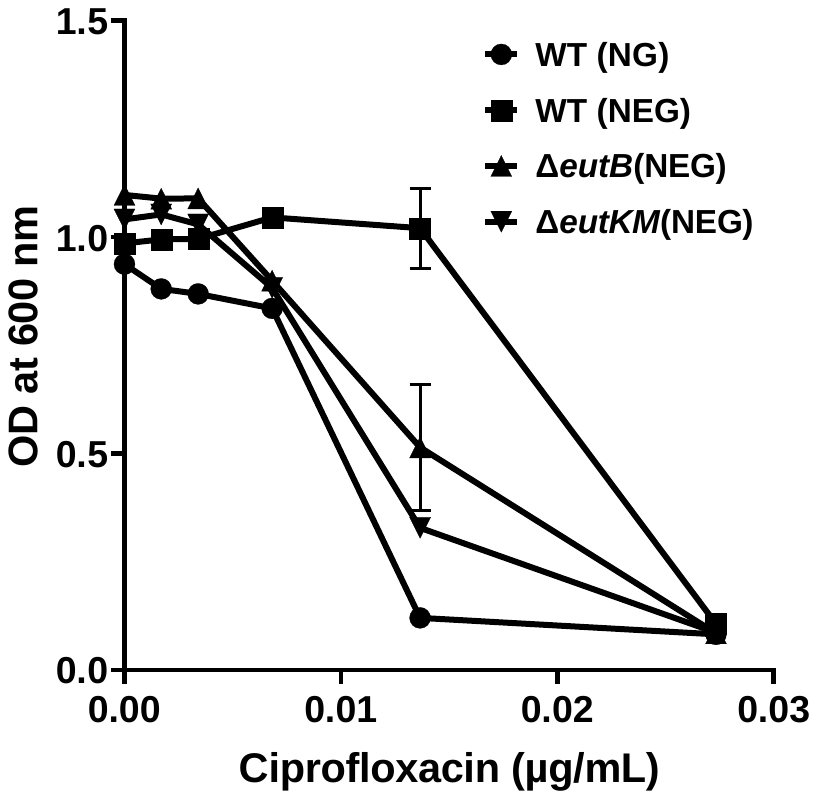

**Figure S3**. The effect of EA utilization in conferring enhanced resistance against ciprofloxacin was shown. The *eutB* and *eutKM* mutants with loss of EA metabolism exhibited null effect. Three biological replicates and four technical replicates were performed. Data is presented as the mean ± SEM.

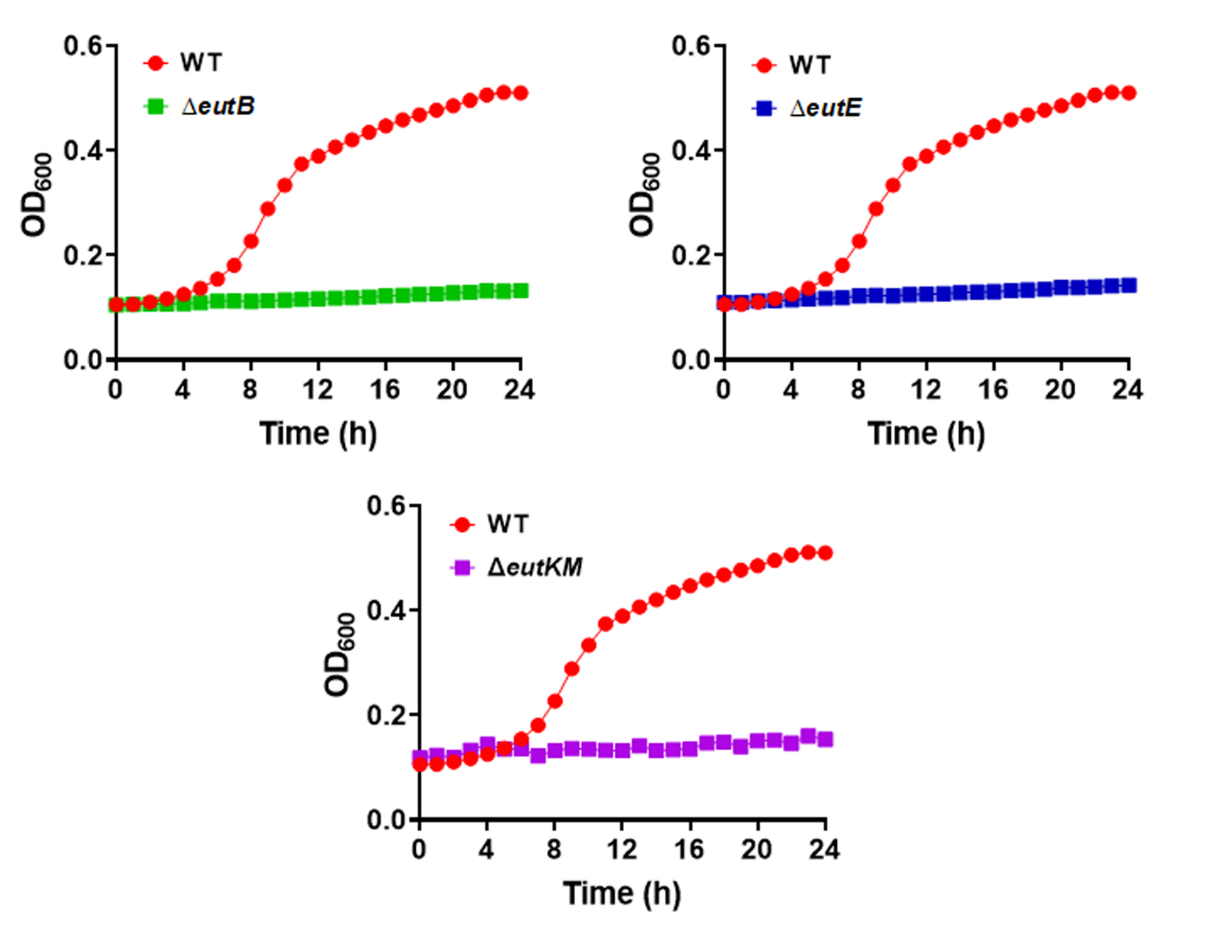

**Figure S4.** Growth curve analysis of WT compared to *eutB, eutE*, and *eutKM* mutants grown in minimal media supplemented with Ethanolamine and Vit. B_12._

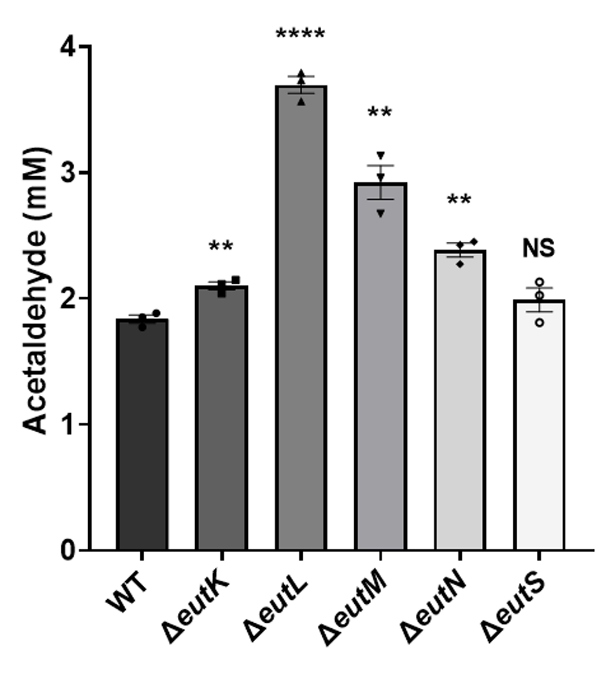

**Figure S5.** Effect of shell protein mutation on retention of acetaldehyde. Acetaldehyde was **r**elease in higher quantity in the shell protein mutants, except an *eutS*. An unpaired *t*-test (two-tailed) was used for statistical significance. NS p > 0.05, *p < 0.05, **p < 0.01, ***p < 0.001, ****p < 0.0001 compared to the wild type control.

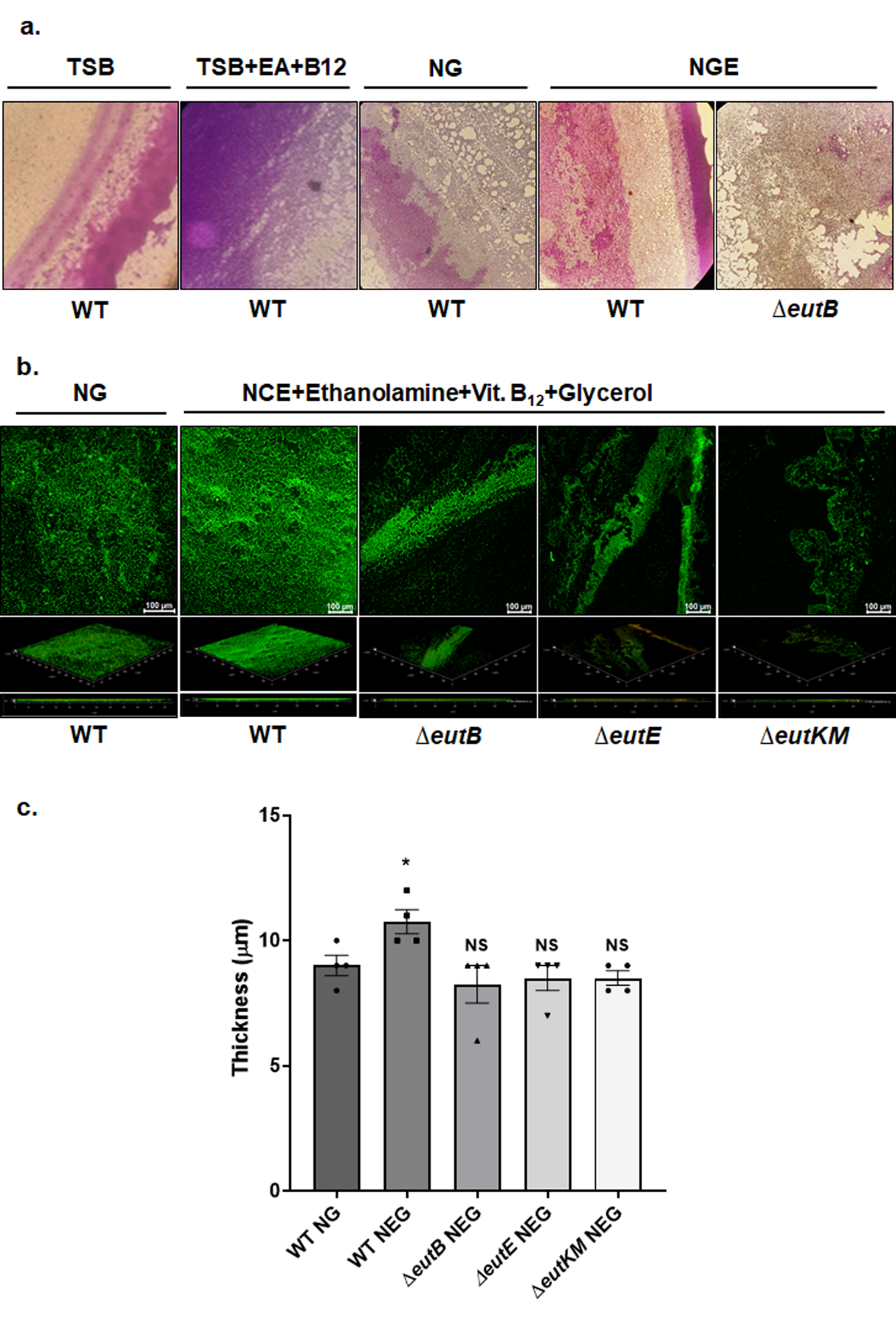

**Figure S6**. Microscopic visualization of biofilm after a) crystal violet staining and b) live/dead staining. Supplementing ethanolamine to rich (TSB) and glycerol media enhances WT *Salmonella*'s biofilm. Biofilm was assessed quantitatively using confocal microscopy with Orthogonal Z-stack and 3D images. c) Thickness measured in 3D images depicted the role of EA metabolism in biofilm formation. Unpaired (Two-tailed) *t*-test indicates a significant increase in biofilm when EA and B_12_ are present in the glycerol media. The mutants with disrupted EA metabolism have no significant effect compared to WT.

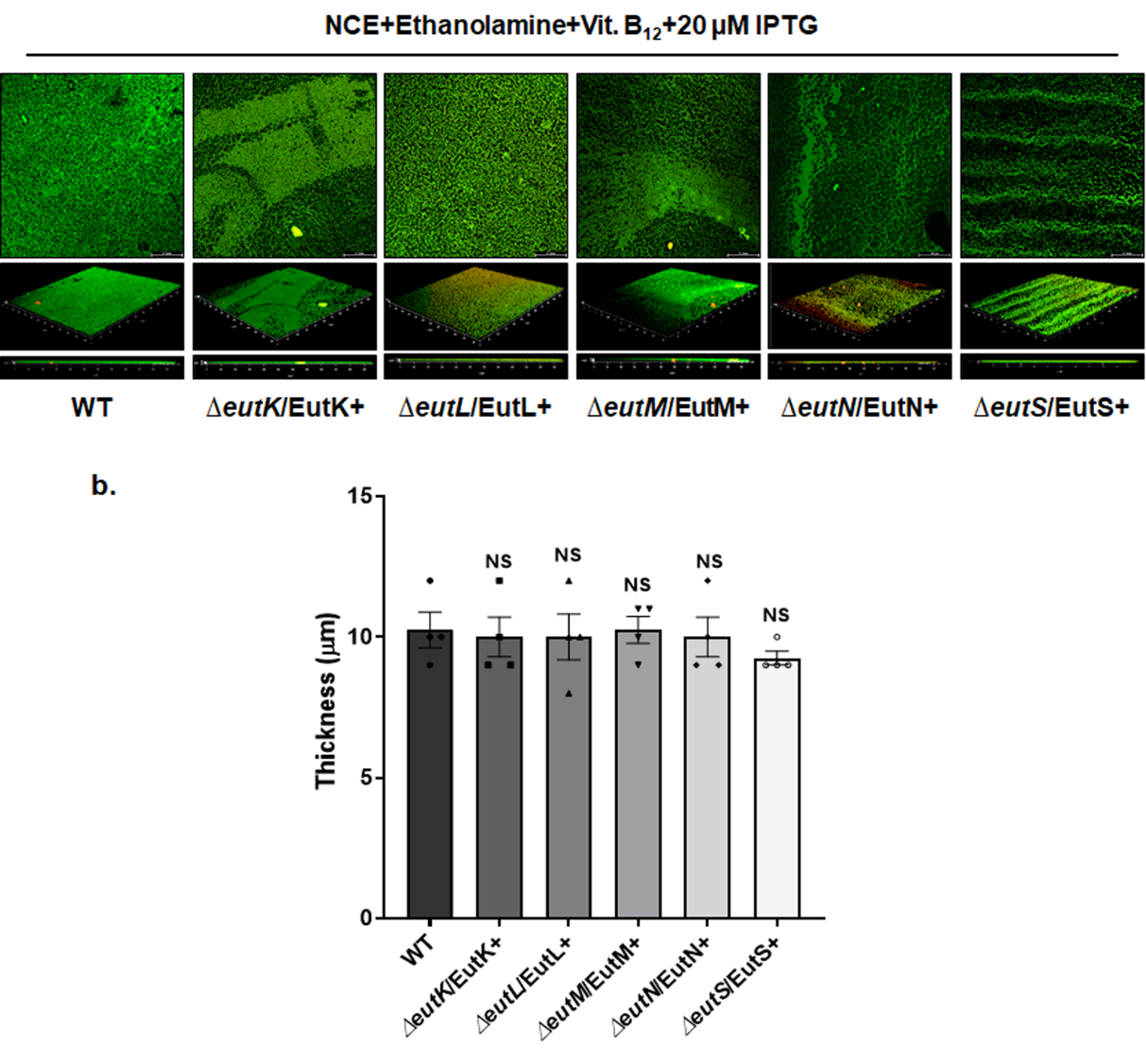

**Figure S7.** Complementation of the mutants with the corresponding genes cloned in pLac22 plasmid. a) Live-Dead Cell Imaging with orthogonal Z-stack and 3D using Confocal Microscopy. b) Thickness of Biofilm measured in 3D images. An unpaired (Two-tailed) *t*-test results in a non-significant difference.

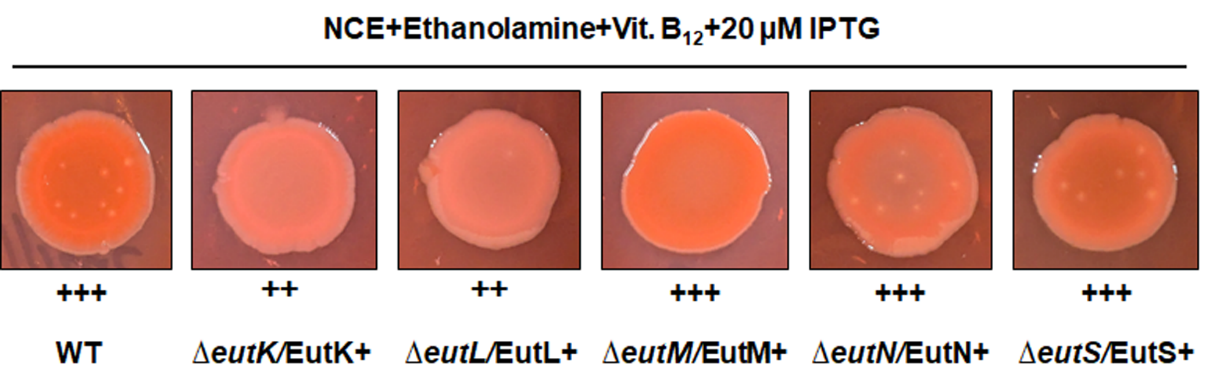

**Figure S8**. Loss of phenotype in Congo red assay were rescued by ectopic expression of the corresponding genes. Expression was carried out by 20 µM IPTG.

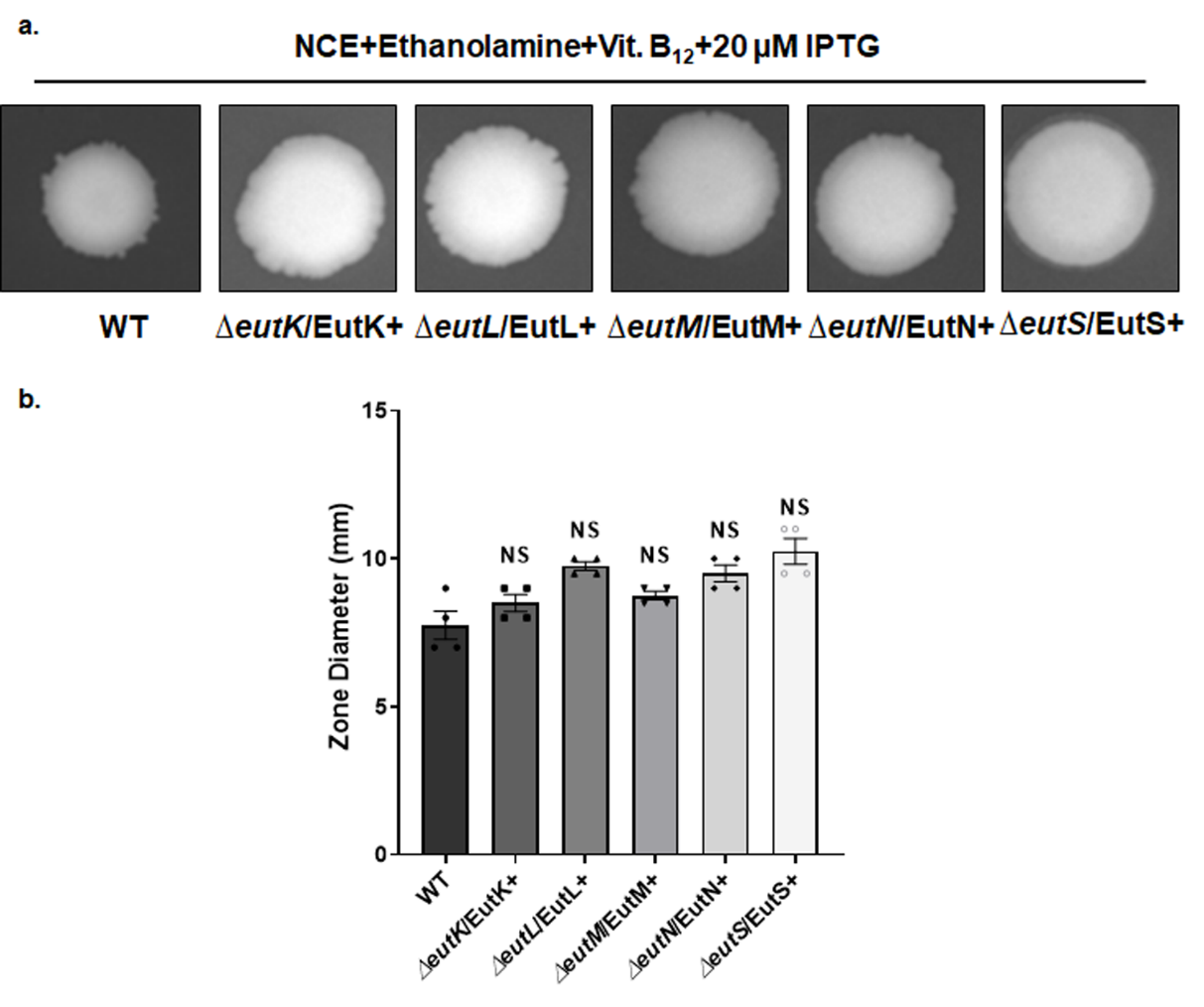

**Figure S9.** Reduced motility in the shell protein mutants was reversed upon complementation with the genes expressed from pLac22 at 20 µM.

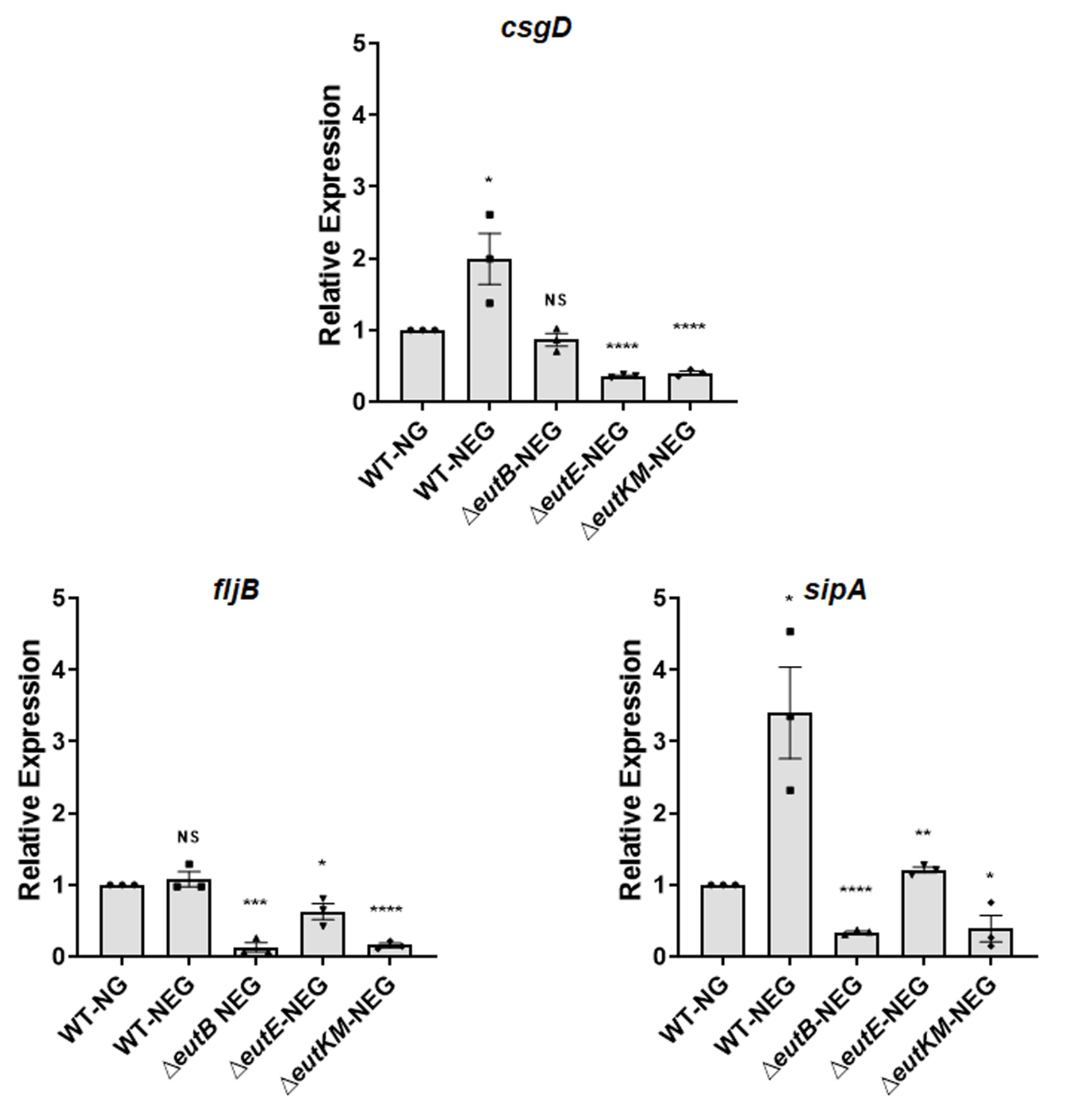

**Figure S10.** Quantitative real-time PCR was carried out for WT supplemented with and without ethanolamine in glycerol media. Gene targets were selected from each experiment group: *csgD* for biofilm, *fljB* for motility, and *sipA* for the cell invasion study. An unpaired *t*-test (two-tailed) was used for statistical significance. NS p > 0.05, *p < 0.05, **p < 0.01, ***p < 0.001, ****p < 0.0001 compared to the wild type control.

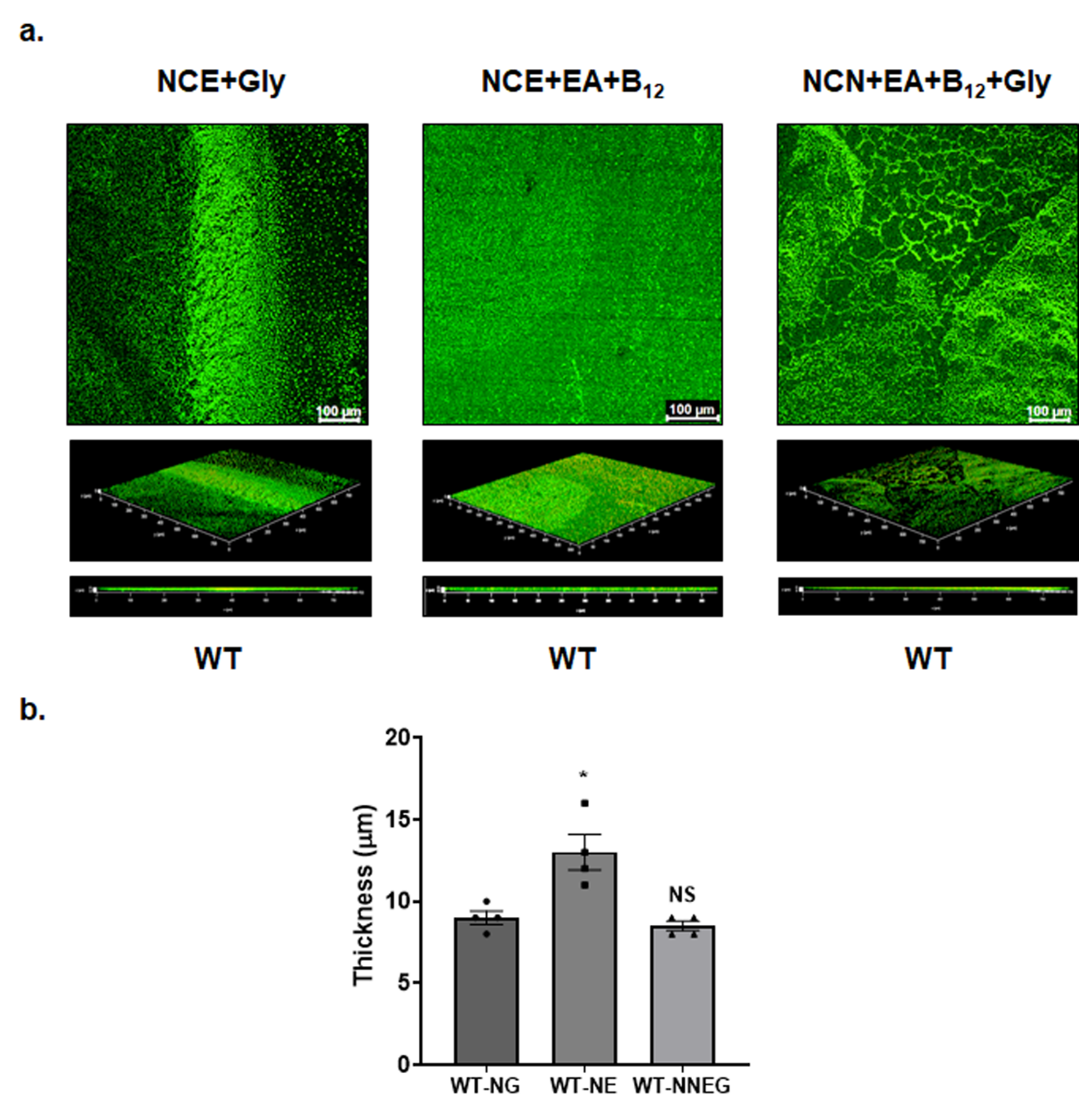

**Figure S11**. Live-Dead Cell Imaging of biofilm with orthogonal Z-stack and 3D using Confocal Microscopy of wild type grown in NCE+Glycerol (NG), NCE+EA+Vit. B_12_ (NE) (EA serving as C source), and NCN+EA+Vit. B_12_+Glycerol (NNEG) (EA serving as N-source). b) Thickness of biofilm measured in 3D images. An unpaired (Two-tailed) *t*-test results in a non-significant difference.

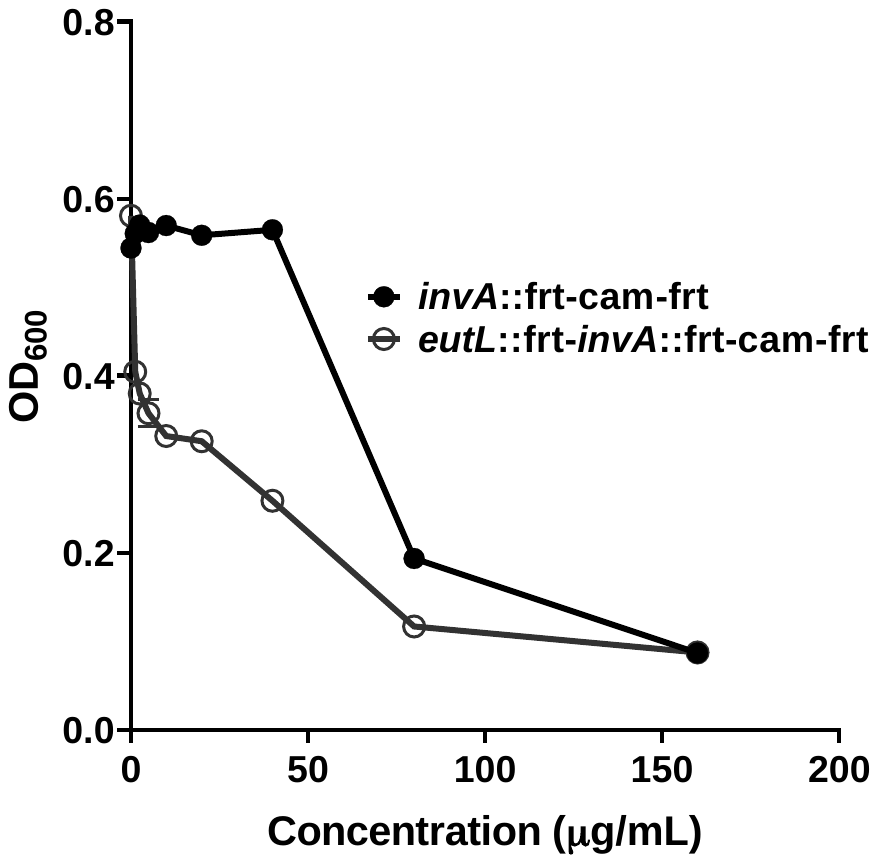

**Figure S12**. Effect of EutL shell protein deletion mutation on the growth of chloramphenicol-resistant *Salmonella*
